# Modification of Seurat v4 for the development of a phase assignment tool able to distinguish between G2 and Mitotic cells

**DOI:** 10.1101/2022.10.27.513990

**Authors:** Steven Watson, Harry Porter, Ian Sudbery, Ruth Thompson

## Abstract

Single cell RNA sequencing (scRNAseq) is a rapidly advancing field which allows for the characterization of the cellular heterogeneity of gene expression profiles within a population. Cell cycle phase is a major contributor to gene expression variance between cells and computational analysis tools have been developed to assign cell cycle phase to scRNAseq datasets. Whilst these tools can be extremely useful, all have the drawback that they classify cells as G1, S or G2/M. Discrete cell phase assignment tool are unable to differentiate between G2 and M and continuous phase assignments tools are unable to identify a region corresponding specifically to mitosis in a pseudo-timeline for continuous assignment along the cell cycle. Bulk RNA sequencing was used to identify differentially expressed genes between mitotic and interphase cells isolated based on phospho-histone H3 expression using fluorescence activated cell sorting. The gene lists were used to develop a Modified Seurat Mitotic Sort (MoSMiS) methodology which can distinguish G2 and M phase cells in single cell RNA sequencing data. The phase assignment tools present in Seurat were modified to allow for cell cycle phase assignment of all stages of the cell cycle identifying a mitotic specific cell population.

## Introduction

Single cell RNA sequencing (scRNA-seq) is a rapidly advancing technology allowing researchers to assay gene expression at a higher resolution than conventional bulk RNA sequencing. This technique has facilitated the characterization of cellular heterogeneity in complex tissues leading to the discovery of novel cell populations. However, these tissues frequently contain cycling cells and cell cycle phase is a major factor driving gene expression variance between cells. This can impact the ability of researchers to define whether two clusters identified in dimensionality reduction plots are two distinct cell types or the same cell type at different cell cycle phases. To address this, many scRNA-seq toolkits apply methodologies to assign cell cycle phase to individual cells in scRNA-seq datasets.

Cell cycle has been recognized as a major contributor to gene expression variance between cells [1–3] leading to the development of various cell cycle computational analysis tools in order to overcome this, including Oscope [4], Peco (Hsiao et al., 2019), reCAT [5] Cyclone [6] and Seurat [7]. Whilst these tools can be extremely useful in the analysis of scRNAseq data for the study of cell cycle specific effects, all of these tools have the drawback that they classify cells as G1, S or G2/M and are unable to differentiate between G2 and M.

Due to the highly condensed nature of mitotic chromatin, it was assumed that transcription was repressed in mitosis [8–10]. This assumption remained in place for many years due to technical limitations, however in the last 5 years, there have been significant advances. In 2017 it was proposed that transcription is maintained at a low level throughout transcription followed by a “wave” of transcription towards the end of mitosis to allow for mitotic exit [11]. Ongoing transcription in mitosis has been observed at centromeres [12] and other chromosome fragile sites [13] and whilst telomeres are thought to be largely transcriptionally silent in mitosis, cells depending on the ALT pathway for telomere maintenance telomere-repeat containing RNA (TERRA) remains associated with telomeres in G2/M [14].

Here we used fluorescent activated cell sorting to separate interphase and mitotic cells prior to RNAseq analysis for the generation of a gene list of mitotically upregulated genes. We then used this list to modify the Seurat cell cycle phase assignment tool to enable differentiation between G2 and M phase cells.

## Methods

### Cell culture

HeLa cells (ATCC) were cultured in DMEM supplemented with 10% FBS at 37°C+ 5% CO_2_. Cells were passaged 1:10 every 3 to 4 days.

### Fluorescence activated cell sorting

Cells were harvested using Trypsin EDTA (Lonza) and washed in phosphate buffered saline (PBS) fixed for 30 minutes at −20°C in 70% Ethanol containing RNaseOUT™ Recombinant Ribonuclease Inhibitor solution (Invitrogen, #10777019). The cells were then washed and rehydrated in PBS before incubation on for 1 hour on ice with Phospho-Histone H3 (Ser10) antibody (Sigma-Aldrich) at 1:1000 in buffer 1 (PBS with 0.5% BSA and 0.25% Triton-X 100). The samples were washed in flow buffer 2 (500 mL PBS, 0.25% Triton-X 100) prior to incubation on ice for 30 mins in the dark with secondary antibody pAb to Rabbit IgG (FITC) (abcam) 4:1000 in buffer 1. Following two washes in PBS, cells were incubated for 30 minutes in 5 mg/ml propidium iodide (Thermo-Fisher Scientific) prior to cell sorting using the FACSMelody (BD Biosciences). Cells were gated based on FITC staining levels and sorted directly into the DNA/RNA lysis buffer.

RNA was extracted using Zymo Research Quick-DNA/RNA Miniprep RNA extraction kit according to the manufacturer’s instructions. RNA quality was assessed using a Samples were run on a Eukaryote total RNA Pico Agilent Bioanalyzer Chip and analysed using Agilent 2100 expert software to calculate RNA integrity numbers (RIN). RNA samples were stored at −80°C and sent to Novogene where total RNA extract was purified to mRNA by poly(A) enrichment. Unstranded paired end sequencing was performed on the Illumina NovaSeq PE150 to a depth of 40 million paired reads and a fragment length of 150 bp generating 12G raw data per sample. 3 biological replicates per condition.

### RNAseq analysis

Quality control of unaligned read data sets was performed using FastQC through the Galaxy Europe GUI [15]. Reads were mapped against the Human Dec 2013 (GRCh38/hg38) (hg38) reference genome using HISAT2 [16]. Samtools were used to check alignment quality. The quantity of reads per gene was calculated using htseq-count [17] against the reference genome. Count matrices were collated and annotated with Ensembl canonical gene symbols and RefSeq IDs. PCA analysis, normalisation and visualisation were all carried out in R.

Differential expression testing was conducted using DEseq2 [18]. Lowly expressed genes and outliers were removed by independent hypothesis filtering of data and Cook’s distance. We selected differentially expressed genes where the adjusted Wald test p-value < 0.001 and ±0.58-fold change, representing 1.5 times more or less gene expression. The genes were separated into an upregulated (Mitotic related) and downregulated (Interphase related) based on positive or negative Log2FoldChange values respectively. Post selecting differentially expressed genes based on adjusted p-value and Log2FoldChange Cutoffs we derived 27 “Mitotic” related genes of interest and 18 “Interphase” related genes of interest.

### Testing cell phase assignment tools

All cell phase assignment protocols were tested using the single cell RNA sequencing count matrix derived from [19] (GSE81682) a murine haematopoietic stem cell/multipotent progenitor single cell RNA sequencing data set and Hu et al., 2019 (GSE129447) using the GSM3713084 HeLa 1 p9 data set.

The Modified Seurat Mitotic Sort uses the cell cycle state assignment approach implemented in Seurat [20] as outlined in the Cell Cycle Scoring and Regression Vignette [21].

The Modified Seurat Mitotic Sort basic stepwise outline is:

1. The count matrix is normalised via a relative count system with an appropriate scale factor the using Seurat NormalizeData function.
2. Variable features are found based on the counts for the marker gene data – S and G2/M in this first instance, using Seurat function FindVariableFeatures
3. Principle component analysis using the scaled and centred counts for the variable S and G2/M marker genes is carried out. This is visualised to verify separation.
4. G2/M, S and G1 phases are assigned, based on G2/M and S phase variability scores using Seurat function CellCycleScoring
5. Steps 2-3 are repeated using cells assigned G2/M phase in step 4 and the list of Interphase and M phase genes.
6. M phase and G2 phases are assigned using a modified CellCycleScoring function. The original function can assign cells to one of three phases using two gene lists. The modified function only assigns to two lists using the list of marker genes provided. (see Extended Code 1).
7. G2 and M assignments are combined with the original G1 and S assignments to full assign all cells to one of G1, S, G2 and M phases.
8. Final phase assignments are then outputted in csv format

After each CellCycleScorng step, assignments and the genes driving these assignments were examined using the DimPlot and RidgePlot Seurat functions respectively.

### K Fold Testing

To ensure that the genes generated in (**Table 1)** where related to directly to interphase or mitotic transcriptome profile cross validation k-fold tests were completed on a range of data sets. Kfold cross validation was tested using the MosMis code on a murine haematopoietic stem cell/multipotent progenitor data set from [19] (GSE81682), a HeLa data set from Hu et al., 2019 (GSE129447) and a Myxoid Liposarcoma data set from Karlsson et al., 2017 (E-MTAB6142).

**Table 1.** Differential Gene Expression Testing via Deseq2. Gene list post statistical significance and expression change filtering. The generated lists gives interphase and mitotic specific gene lists, thus allowing for the customisation and optimisation of the Seurat CellCycleScoring pipeline. Canon gene symbols are used unless unnamed/novel in which case the Ensembl number is provided. The gene list was generated post filtration of regulated expression levels (log2FoldChange < 0.58) and statistically significant differential expression (padj < 0.001).

| Downregulated Genes (Interphase) | Upregulated Genes (Mitotic) |
| --- | --- |
| E2F1 | CENPE |
| CCNE1 | KNL1 |
| FBXL20 | PIMREG |
| DTL | PLK1 |
| ENSG00000273759 | KIF14 |
| RMI2 | TPX2 |
| ZMYND19 | KIF20A |
| MCM5 | SAPCD2 |
| ZNF367 | KNSTRN |
| FRAT1 | PRR11 |
| BRD2 | NUF2 |
| ENSG00000272106 | ASPM |
| PPP1R3C | CEP55 |
| ENSG00000275484 | BUB1 |
| UNG | SGO2 |
| IFI27L1 | GAS2L3 |
| CDC6 | NEK2 |
| MIR25 | HMMR |
|  | DEPDC1 |
|  | DLGAP5 |
|  | ARL6IP1 |
|  | NUSAP1 |
|  | CCNA2 |
|  | VANGL1 |
|  | CDC20 |
|  | KIF4A |
|  | KIF20B |

Datasets were transformed using rlog or vst transformation and analysed similarly to the above outlined process except that 25% of either the interphase or mitotic generated gene list of interest were removed first. This was performed 4 times for each 25% of the gene population of the interphase and mitotic genes. The expression of the removed genes in each of the 4 tests (Interphase and Mitotic removal were tested separately) were then analysed for overall net expression in the populations which had been phase assigned by the remaining 75%.

### RT-qPCR

Mitotic and interphase cells were separated by mitotic shake off and RNA was extracted using the Zymo RNA extraction kit according to the manufacturer’s instructions. cDNA was generated using an RNA to cDNA kit (Thermo Fisher Scientific). cDNA was mixed with the specified primers and SYBR green PCR master mix (ThermoFisher scientific) and 40 cycles of real time PCR was carried out on a QuantStudio 7 Pro Real-Time PCR System (Thermo Fisher Scientific). We performed 3 biological replicates of each condition, and qPCR was carried out for each of these in 3 technical replicates. Data was analysed using the ddCT method (Livak, 2001) normalised to the geometric mean of two housekeeping genes: 18S and HPRT1. Statistical analysis was conducted by a paired t test, the mitotic enriched samples compared to corresponding control. Data analysed via Prism.

### Gene Function Orthologies

The gene lists generated by our RNAseq analysis of mitotic vs interphase cells were analysed by GOrilla for gene ortholog. The total gene list from the bulk RNAseq count matrix was used as the background gene list. The “interphase” and “mitotic” related genes of interest as seen in table 1, used as the gene sets for function GO analysis. A P-value threshold of 10^−3^ selected for testing. Significant results scored in 10^−3^ to 10^−5^, 10^−5^ to 10^−7^, 10^−7^ to 10^−9^ and 10^−9^ and below p value bands used to determine levels of function significance.

### Data and Code Availability

The data for testing was accessed from the gene expression omnibus. The HeLa cell dataset is available under accession code GSE129447 and the murine progenitor dataset is available under the code GSE81682.

The completed Modified Seurat Mitotic Sort (MosMis) code is available in supplemental data.

## Results

### Generation of a list of genes upregulated in mitotic cells

We tested several phase assignment tools before settling on Seurat for modification. Here we show cell cycle analysis using both a discrete (cyclone, **Figure 1A**) and a continuous (peco **Figure 1B**) phase assignment tool. Using an scRNAseq dataset of HeLa cells at an early passage number (GSE129447) GSM3713084 [22] we found that Cyclone [6] did not provide sufficient ability to subset out mitotic cells **(Figure 1A).** Each singular point in the analysis represents a single cell tested, the default Cyclone function showing clearly cells scoring highly as G1 or G2/M. We were unable to attain clear clustering of the dataset.

**Figure 1.**
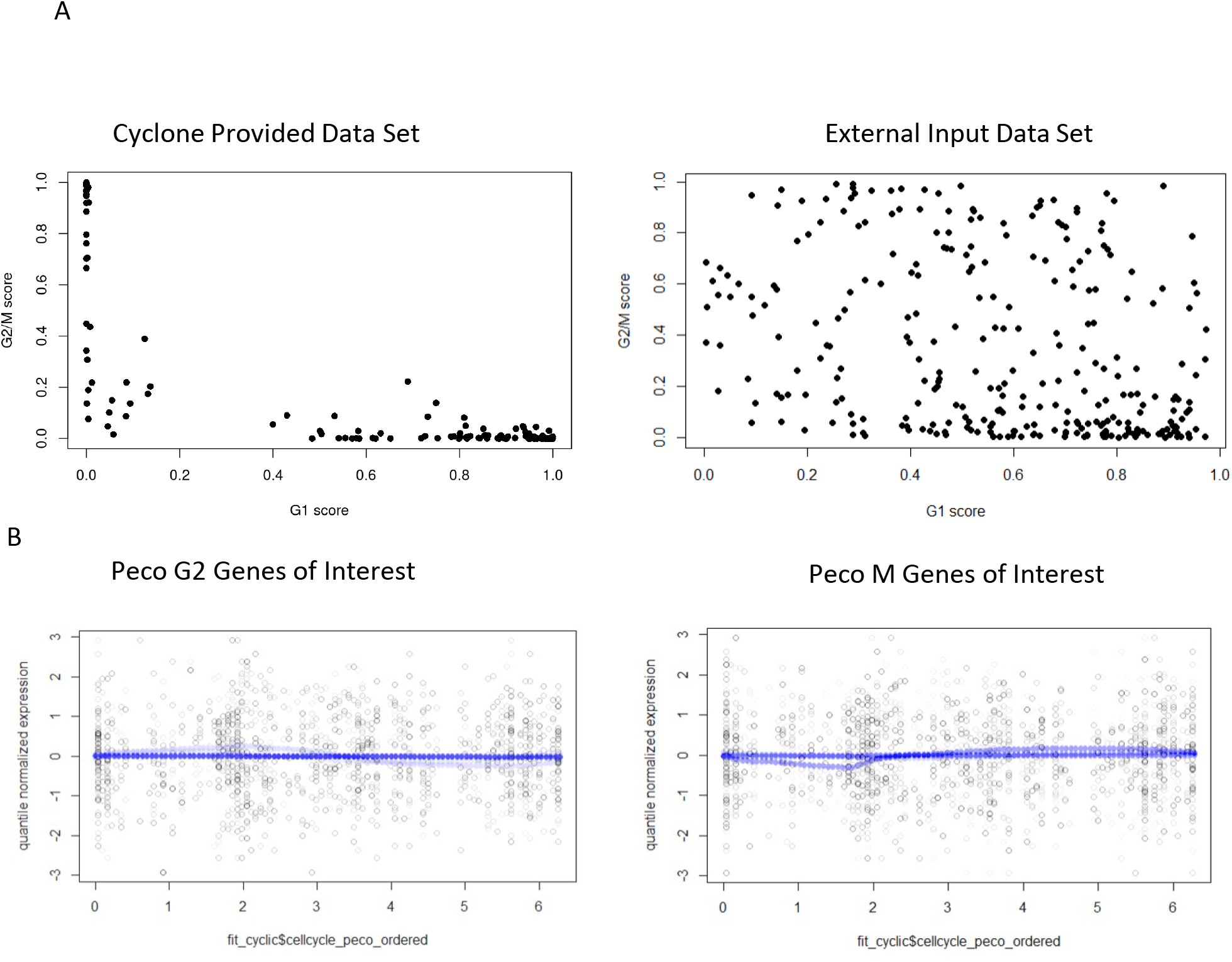
Finding a phase sorting code base to analyse and detect mitotic cells. A) Dataset GSE129447 GSM3713084 was tested and graphed in R, using Cyclone a subfunction of scran (https://bioconductor.org/packages/release/bioc/html/scran.html B) Same dataset GSE129447 GSM3713084 testing and graphed in R, using peco (http://www.bioconductor.org/packages/release/bioc/html/peco.html

Peco (Hsiao et al., 2019) orders cells along the cell cycle and estimates the cyclic trend of a given gene based on FUCCI phase and plotting per cell expression levels for the assigned gene giving a pi value for each (white and black dots). Peco then generates a normalised expression trend of the gene (blue line). When testing a range G2 and M phase specific genes the normalised expression line became smoothed across both the G2 and M gene tested and we were unable to sufficiently generate points of differentiation between G2 and M phase aligned genes that fit the 101 gene used as cell phase predictors in Peco. **(Figure 1B).** We found no distinguishing peaks or troughs in the quartile normalised expression or the cyclic fitted genes of interest when using Peco.

Seurat, an open-source toolkit for analysis of scRNA-seq data, was selected for modification as its cell cycle phase prediction methodology can be adapted by providing a manual gene list rather than using the default gene list provided within the package. The cell cycle prediction function of the Seurat tool separates G1, S and G2/M cell populations. However, it can use a supplied gene list instead of the default list to underlie its cell cycle state assignment. We reasoned that if supplied with a list of G2 and M specific genes, the same approach may be able to further separate G2 from M cells.

To modify the Seurat for assignment of a mitotic fraction, a list of genes upregulated in mitotic cells was required. To generate this list, we stained cells using an antibody to histone H3 phosphorylated at Serine 10 (pH3) as a marker of mitosis and sorted mitotic cells using fluorescence activate flow cytometry (**Figure 2).** We were able to cleanly separate the pH3 expressing population from the non-pH3 expressing population and confirmed that the pH3 positive population all contained 4N DNA content (**Figure 2B**). RNAseq analysis revealed clear differentially expressed gene profiles between the mitotic and interphase populations (**Figure 3A).** At a standard threshold for differential expression (adjusted p-value < 0.05), we identified 76 genes preferentially expressed in interphase and 86 preferentially expressed in mitosis. To identify a high confidence set of phase specific genes, we used a more restrictive adjusted p-value cut-off of ≥0.001 **(Figure 3B).** The gene lists shown in table 1 are the final lists used in our modified Seurat mitotic sort code for the separation of mitotic and interphase populations of the G2/M population (**Table 1**). To confirm upregulation of the “mitotic” genes in mitotic cells, we used qRT-PCR to measure the expression of the 4 highly upregulated “mitotic” genes, in interphase and mitotic HeLa cells separated by mitotic shake off, an orthogonal method of isolating mitotic cells. All four of the mitotic genes selected showed significantly higher expression in the mitotic fraction than the interphase fraction **(Figure 4A)**.

**Figure 2.**
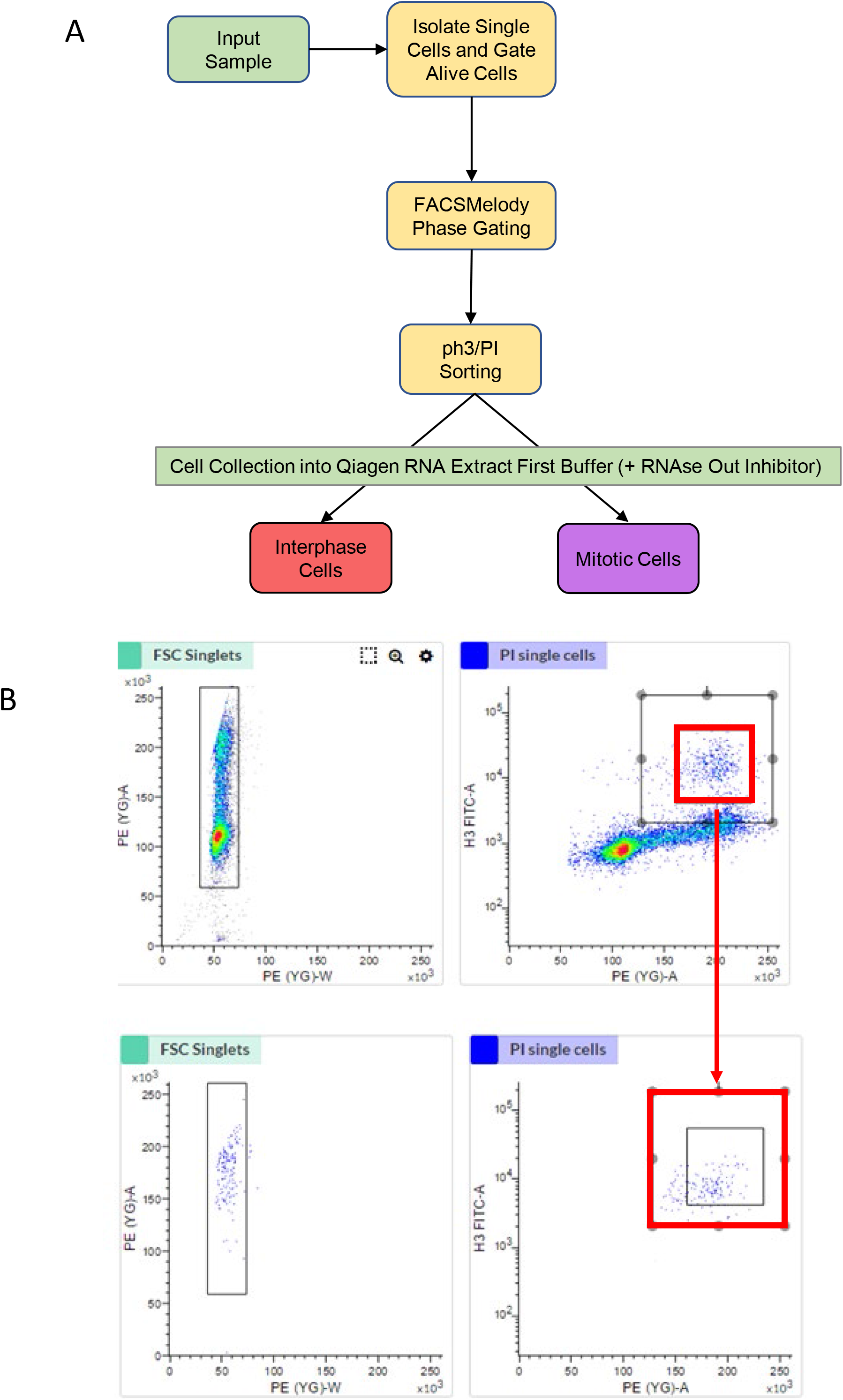
Separation of interphase and mitotic cells. A) Workflow for separation of mitotic and interphase cells using the FACSMelody. Cells were sorted based on phospho-histone H3-FITC staining. **B)** HeLa cells were stained for pH3-FITC and PI and seprataed using the FACSMelody based on FITC staining. Representative plots showing separation of the phospho-histone H3 positive population.

**Figure 3.**
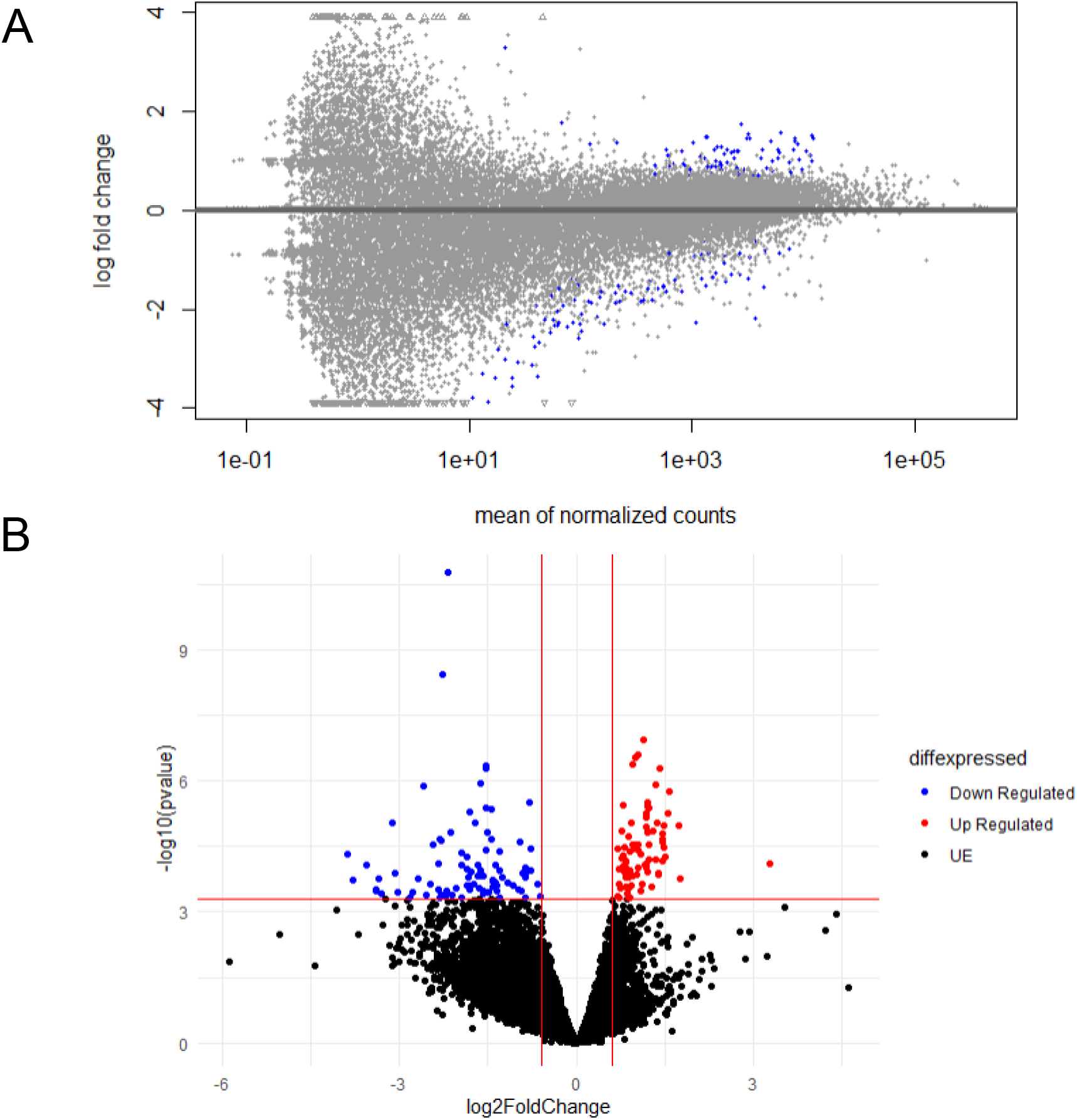
Differential Gene Expression Testing via Deseq2. A) MA plot of RNA-seq data. Scatter plot of log2 fold changes versus the mean of normalized counts. Each dot represents a singular gene across all present, blue dots represent differentially expressed genes under the preset padj<0.05 threshold. Grey dots are not statistically significant. The differentially expressed genes blue points with a positive log fold change are indicative of mitotic related genes. The differentially expressed genes blue points with a negative log fold change are indicative of interphase related genes. Data was plotted in R using plotMA from deseq2 dds values regressing out the effect of treatment and replicate. B) Differentially expressed genes in RNA-seq volcano plot. Red data points represent upregulated mitotic weighted genes, blue points represent downregulated interphase weighted genes. log2Foldchange of ±0.58 and padj value of ≤ 0.05.The volcano plot was created using the count matrix input and graphed via ggplot2. X-axis is log2 fold change, y-axis is statistical significance −log10 (adjusted p-value).

**Figure 4.**
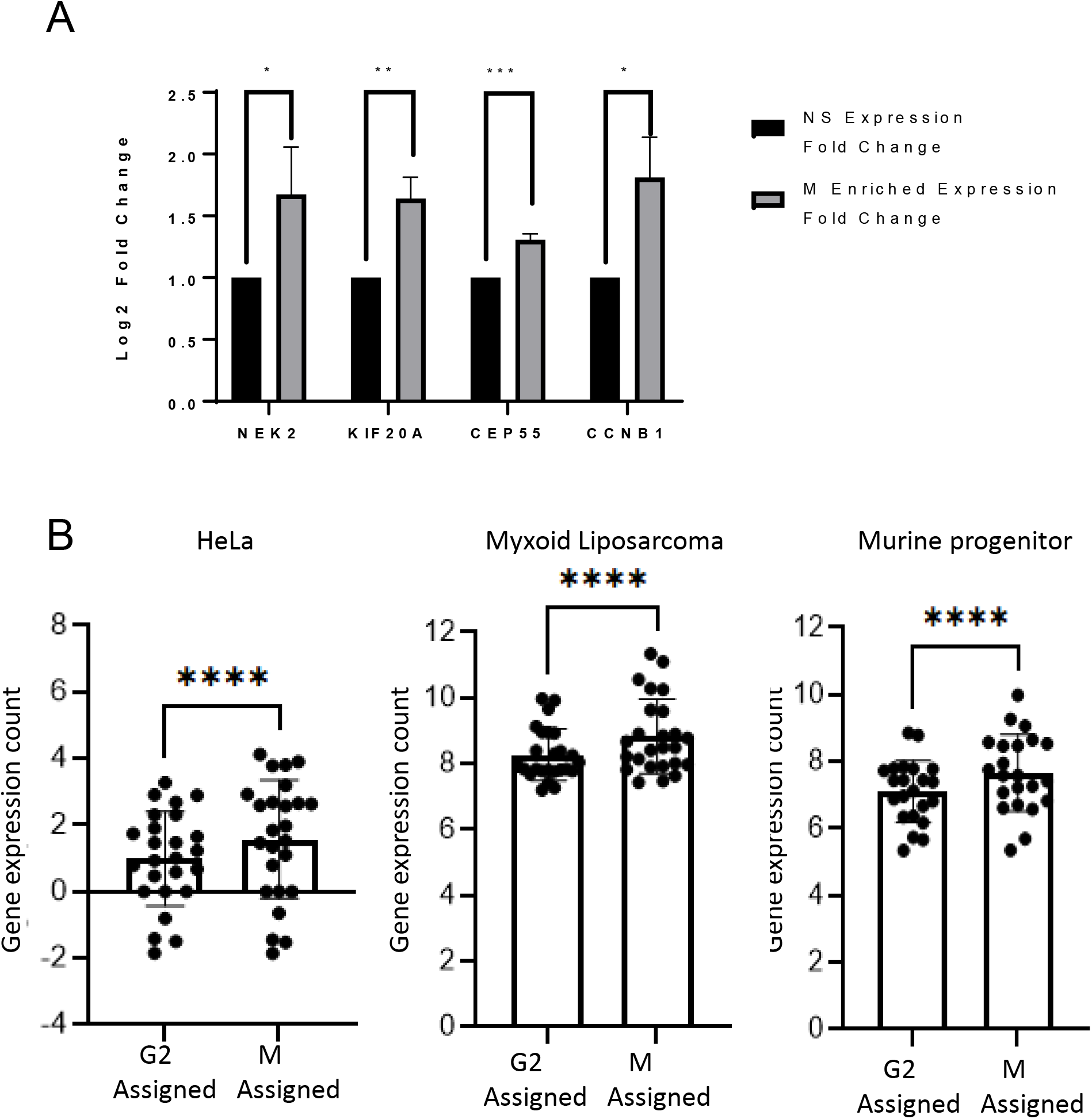
Validation of mitotic genes list. *A)* qRTPCR was carried out on cDNA from RNA extracted from interphase (attached) or mitotic (shaken-off) HeLa cells. B) Four fold K test of gene exclusion list expression levels on log2 count matrix. Paired t test performed compared to corresponding control. * = P ≤ 0.05, ** = P ≤ 0.01, *** = P ≤ 0.001, ****= P ≤ 0.0001. Test datasets used GSE129447-1, E-MTAB6142, GSE81682. Tests were completed in R using a looped Modified Seurat Mitotic Sort designed to test with a 25% excluded mitotic or interphase gene list each time. N=4.

Testing function gene orthologies revealed that the most significantly enriched functional pathway in our “mitotic phase” genes was “microtubule, tubulin binding or other cytoskeletal protein binding”. Other terms enriched in this set involved the kinetochore and anaphase promoting complex and ATPase and microtubule motor proteins **(Table 2).** We were therefore confident that this gene list represented set of genes whose expression was higher in mitosis than in interphase and might make a suitable set of markers for identifying mitotic cells.

**Table 2.** Go Function links for Mitotic related genes of interest. Microtubule, tubulin binding or other cytoskeletal protein binding all logically link a gene list related to mitotic function and expected roles in mitosis.

|  |  |  |
| --- | --- | --- |
| Mitotic Gene of Interest Grouped via GO Function |  |  |
| GO Linked Function | Ensembl | Gene Symbol |
| Anaphase-Promoting Complex Binding | ENSG00000117399. | CDC20. |
| ATP Binding | ENSG00000112984.<br>ENSG00000138182.<br>ENSG00000090889.<br>ENSG00000166851.<br>ENSG00000169679.<br>ENSG00000117650. | KIF20A. KIF20B. KIF4A. PLK1. BUB1. NEK2. |
| Kinetochore Protein | ENSG00000137812. | KNL1. |
| Microtubule Motor Protein | ENSG00000112984.<br>ENSG00000138182.<br>ENSG00000090889. | KIF20A. KIF20B. KIF4A. |
| Microtubule/ tubulin Binding | ENSG00000118193.<br>ENSG00000112984.<br>ENSG00000138182.<br>ENSG00000090889.<br>ENSG00000128944.<br>ENSG00000137804.<br>ENSG00000166851.<br>ENSG00000088325.<br>ENSG00000138778.<br>ENSG00000126787.<br>ENSG00000139354. | KIF14. KIF20A. KIF20B. KIF4A. KNSTRN.<br>NUSAP1. PLK1. TPX2. CENPE. DLGAP5.<br>GAS2L3. |
| Mitotic Kinase | ENSG00000166851. | PLK1. |
| Protein Binding | ENSG00000170540.<br>ENSG00000066279.<br>ENSG00000145386.<br>ENSG00000138180.<br>ENSG00000024526.<br>ENSG00000072571.<br>ENSG00000186193.<br>ENSG00000163535.<br>ENSG00000173218. | ARL6IP1. ASPM. CCNA2. CEP55. DEPDC1.<br>HMMR. SAPCD2. SGO2. VANGL1. |

### Using MoSMiS to phase assign scRNAseq datasets

We used our list of mitotically expressed genes to separate a previously assigned G2/M population of cells into G2 and M sets. To test the robustness of these assignments, we used 4-fold validation. The robustness of the mitotic separation was tested by dividing the mitotic gene list into four folds, for each fold we separated G2 from M cells using 75% of the genes in our gene list, holding out 25% of the genes. We then tested the separation by examining the expression the held out 25%. This was carried out 4 times, each time excluding a different selection of genes until all genes had been tested. In all four folds, the excluded genes consistently showed higher expression in the cells which had been designated as mitotic by the remaining genes. This was conducted using different scRNAseq datasets and each showed the same (**Figure 4B**).

We can now demonstrate phase assignment of an scRNAseq dataset GSE81682. The first step of the analysis is the original Seurat code (**Figure 5A**) followed by a secondary sort of the G2/M assigned population into distinctly grouped G2 and Mitotic populations (**Figure 5B**). This dataset was used in the Seurat vignette [23] with their numbers shown in **Figure 5C (Column 1**). We first replicated their phase assignment by running it through the unaltered Seurat code in R as shown in **Figure 5C** (Column 2). Our data were within a few percent of theirs for each phase of the cell cycle. Finally, **figure 5C** (Column 3) shows the same dataset analysed for cell cycle phase assignment by MoSMiS. Whilst as expected, the G1 and S phase values do not change, the G2/ M population is now cleanly separated into two distinct G2 and M populations.

**Figure 5.**
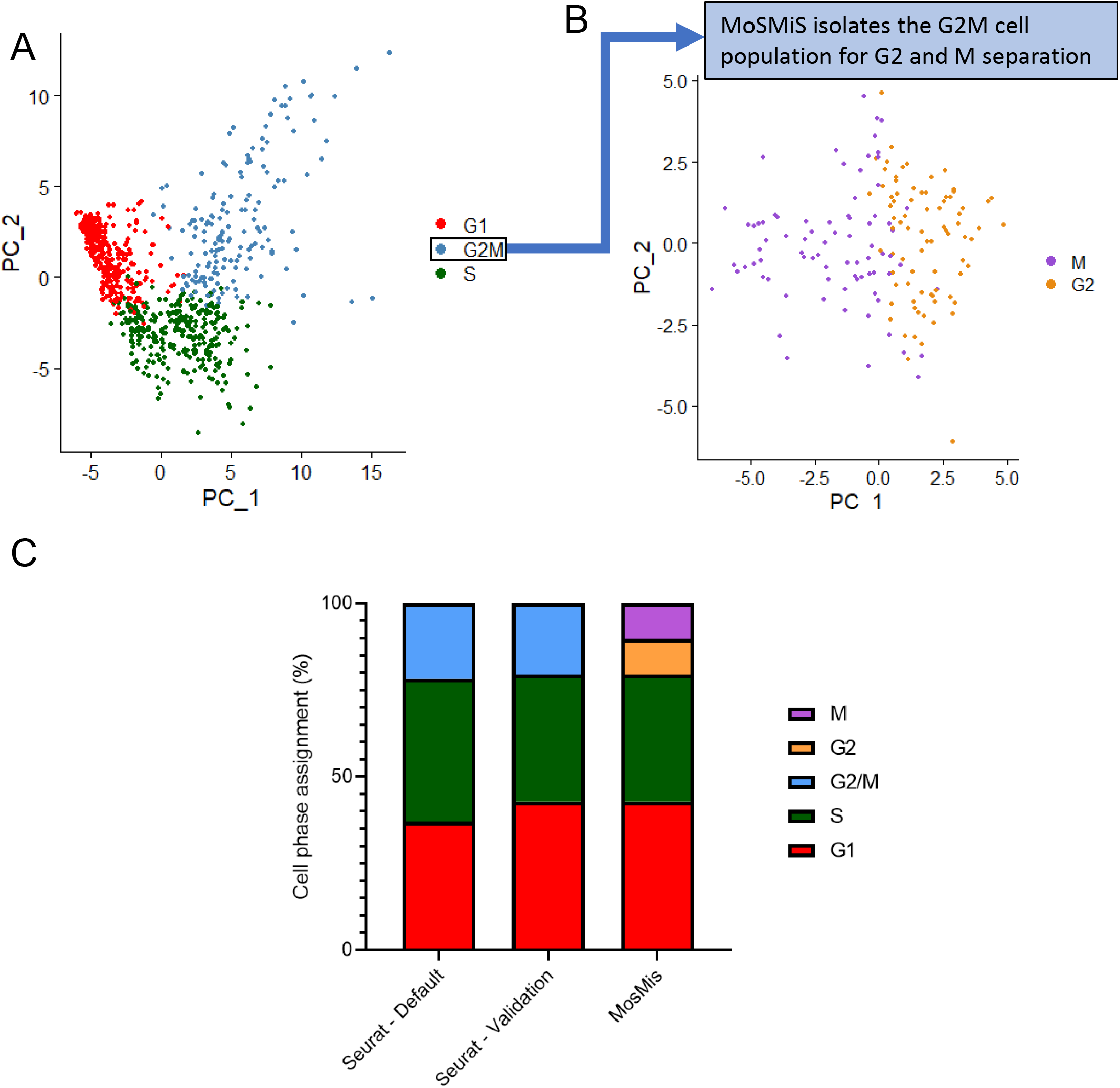
Comparative phase assignment from default Seurat Cell Cycle Sorting and Modified Seurat Mitotic Sort. Dataset GSE81682 (murine progenitor cells) was used as the test data set as this was presented with the Seurat vignette. A) Shows the initial step function of Seurat based cells using RC normalization. Sorting is given in the distributed percentage of cell cycle phase assignments across the input data seen in both the plotted per cell PCA data and the total cell phase percentages. PCA shown is calculated from all input cells from GSE81682 using the Suerat G2/M and S separation gene of interest sets B) MoSMiS add a stage of secondary Seurat Cell Cycle Scoring on the isolated G2/M cell population. PCA calculated using the generated G2 and M specific separation genes of interest set on the subsetted G2/M cells assigned in A. C) Shows the cell cycle assignment as dictated by the Seurat vignette. Tests were completed in R using the Modified Seurat Mitotic Sort (MoSMiS) against default Seurat phase assignment.

## Discussion

Using fluorescence activated cell sorting of HeLa cells we were able to separate phospho-histone H3 positive cells from phospho-histone H3 negative cells. Histone H3 is phosphorylated on ser 10 in mitosis correlating directly with chromosome condensation [24,25]. The pH3 marker has been shown to be specific for mitotic cells with good reproducibility in breast cancer cells [26].

For validation of our gene lists, mitotic and interphase cells were separated using the mitotic shake-off technique to provide an alternative method of separation as pHH3 does not always exactly correlate with mitosis [27,28]. When animal cells enter mitosis the focal complexes which anchor them to their surroundings are disassembled [29,30] and they round up to become spherical meaning they can be more easily detached from a culture plate. We were able to see significantly increased expression levels of the top 4 genes from the mitotic list in the shaken-off population as compared to the attached population confirming upregulation in mitosis.

For further validation of our list, 25% of genes were excluded from the list and cells were phase assigned based on the remaining 75% to test the robustness of the MosMis phase assignment. This was repeated 4 times with a different subset excluded each time. We found that in three different cell lines (two human cancer and one mouse progenitor cell line) the phase assignment agreement of assigned cells remained similar between subsetting tests. The gene phase specific expression levels showing significantly higher expression levels of the excluded mitotic genes in the cells assigned as mitotic using the remaining 75% of genes, therefore highlighting the “mitotic” related GOI list ability to accurately assign mitotic specific genes when used with MosMis regardless of cell line tested.

Finally, we phase assigned the murine progenitor data set [19] using MoSMiS as this same dataset was used as an example in the Seurat vignette [23] which could be used as a comparison. As expected, we found we were able to closely replicate the phase assignment of this dataset demonstrated by the vignette [23]. Furthermore, PCA analysis of the separated G2M population showed clear clustering for the G2 and M phase assigned cells.

In summary we have modified Seurat v4 [20] to develop a tool for cell cycle phase assignment which is able to distinguish between G2 and M phase cells allowing for greater depths of analysis of single cell RNA seq datasets going forward.

## Supporting information

Code Markdown

