## Supplementary material for "Modification of Seurat v4 for the development of a phase assignment tool able to distinguish between G2 and Mitotic cells": Code Markdown

### MosMis

Steven Watson

26/10/2022

#### Input packages required

Note this version used Ensembl gene inputs needs conversion from canon gene names to Ensembl assignment numbers to function

```
library(ggplot2)
library(readr)
library(dplyr)
library(tidyr)
library(devtools)
library(SingleCellExperiment)
library(doParallel)
library(foreach)
library(Seurat)
library(plyr)
library(tibble)
```

#### Test scRNA seq data set and naming structure

Count matrix is input we demonstrate this the data set from murine hematopoietic progenitors in the appropriate format (Nestorowa et al., Blood 2016). Data set used can be found at https://www.ncbi.nlm.nih.gov/geo/query/acc.cgi?acc=GSE81682.

```
#Assign GSE or identifier - Will write with this prefix on output files
GSE <- "Nestorawa_"

#Read in count data
read_data <- read.csv("data/nestorawa_forcellcycle_expressionMatrix_Ensembl.csv", header = TRUE,
    as.is = TRUE , row.names = 1)

#Assigns column numbers for npcs used in RunPCA Functions
npcs_num <- ncol(read_data)
if (npcs_num  < 50) {
npcs_use <- npcs_num-1
} else {
npcs_use <- 50
}
```

#### Intial G1, S and G2/M Seurat cell phase assignment

We intial assign the input count matrix G1, S and G2/M phase assignment using the Seurat Cell Cycle Scoring function with relative count normalization. Seurat functions derived from https://satijalab.org/seurat/articles/cell_cycle_vignette.html.

Chosen input in this step uses the default seurat provided s and G2M genes to isolate a G2M only fraction using Ensembl Seurat Generic Input - converted from cc.genes.updated.20.

```
cell_marker_suerat <- read_csv("data/Seurat Genes Ensembl.csv")
seurat.s.genes <- cell_marker_suerat$seurat.s.genes
seurat.g2m.genes <- cell_marker_suerat$seurat.g2m.genes
seurat.s.genes <- na.omit(seurat.s.genes)
seurat.g2m.genes <- na.omit(seurat.g2m.genes)
```

A Seurat object is created from raw data

```
#Create a Seurat object from raw data
rnaseq_data_seuratgenes <- CreateSeuratObject(counts = read_data)
```

Normalisation step of count matrix using Relative counts, input G2M and S feature counts for each cell are divided bt total counts and scaled by scale.factor. RC uses no log transformation, in further steps will log G2M fraction do not want to repeat log function.

```
rnaseq_data_var_seuratgenes <- NormalizeData(
  rnaseq_data_seuratgenes,
  assay = "RNA",
  normalization.method = "RC",
  scale.factor = 10000,
  margin = 1,
  verbose = TRUE
)
```

A mean variability plot is used to assign outliers in data set, the selection method vst was chosen. Vst fits the relationship of log(variance) and log(mean) using local polynomial regression (loess). The function then standardizes the feature valued using the mean and expected variance from that fitted relationship. The standardized values are used to calculate the feature variance.

```
rnaseq_data_var_seuratgenes <- FindVariableFeatures(rnaseq_data_var_seuratgenes, selection.method = "vs
```

Features are centered and scaled in the dataset.

```
rnaseq_data_var_seuratgenes <- ScaleData(rnaseq_data_var_seuratgenes, features = rownames(rnaseq_data_s
```

```
## Centering and scaling data matrix
```

Post scaling a PCA dimensionality reduction is run. Uses npcs assigned at chunk start and uses variable features of input count matrix with default seurat gene list of interest. PrintPCAParams can be run for more detail.

```r
rnaseq_data_var_seuratgenes <- RunPCA(rnaseq_data_var_seuratgenes, features = VariableFeatures(rnaseq_da
```

```
## PC_ 1
## Positive:  ENSG00000111348, ENSG00000089327, ENSG00000102879, ENSG00000162511, ENSG00000067225, ENSG
## Negative:  ENSG00000167741, ENSG00000164932, ENSG00000164010, ENSG00000111843, ENSG00000107789, ENSG
## PC_ 2
## Positive:  ENSG00000197694, ENSG00000005381, ENSG00000085491, ENSG00000140471, ENSG00000138772, ENSG
## Negative:  ENSG00000205639, ENSG00000130203, ENSG00000172794, ENSG00000179348, ENSG00000169062, ENSG
## PC_ 3
## Positive:  ENSG00000124216, ENSG00000130592, ENSG00000180758, ENSG00000159579, ENSG00000168811, ENSG
## Negative:  ENSG00000105374, ENSG00000130203, ENSG00000179348, ENSG00000149516, ENSG00000182264, ENSG
## PC_ 4
## Positive:  ENSG00000066739, ENSG00000071575, ENSG00000178982, ENSG00000161011, ENSG00000169062, ENSG
## Negative:  ENSG00000137804, ENSG00000175063, ENSG00000185156, ENSG00000131747, ENSG00000141076, ENSG
## PC_ 5
## Positive:  ENSG00000170540, ENSG00000129170, ENSG00000138764, ENSG00000159625, ENSG00000175063, ENSG
## Negative:  ENSG00000117400, ENSG00000146281, ENSG00000108924, ENSG00000149564, ENSG00000221852, ENSG
## PC_ 6
## Positive:  ENSG00000081189, ENSG00000177272, ENSG00000105583, ENSG00000186766, ENSG00000172382, ENSG
## Negative:  ENSG00000140470, ENSG00000140287, ENSG00000101000, ENSG00000149516, ENSG00000033170, ENSG
## PC_ 7
## Positive:  ENSG00000030419, ENSG00000033170, ENSG00000112977, ENSG00000139193, ENSG00000174059, ENSG
## Negative:  ENSG00000105583, ENSG00000172382, ENSG00000005961, ENSG00000081026, ENSG00000136449, ENSG
## PC_ 8
## Positive:  ENSG00000136449, ENSG00000110375, ENSG00000182578, ENSG00000112799, ENSG00000152804, ENSG
## Negative:  ENSG00000172543, ENSG00000115738, ENSG00000112081, ENSG00000184402, ENSG00000186895, ENSG
## PC_ 9
## Positive:  ENSG00000174059, ENSG00000112977, ENSG00000119535, ENSG00000174944, ENSG00000159579, ENSG
## Negative:  ENSG00000146281, ENSG00000129757, ENSG00000101000, ENSG00000136449, ENSG00000100385, ENSG
## PC_ 10
## Positive:  ENSG00000019582, ENSG00000160781, ENSG00000197561, ENSG00000143153, ENSG00000149516, ENSG
## Negative:  ENSG00000127481, ENSG00000136449, ENSG00000188404, ENSG00000110375, ENSG00000167850, ENSG
```

A heat map of the points of variance PC1 to PC10 are plotted

```r
DimHeatmap(rnaseq_data_var_seuratgenes, dims = c(1:10))
```

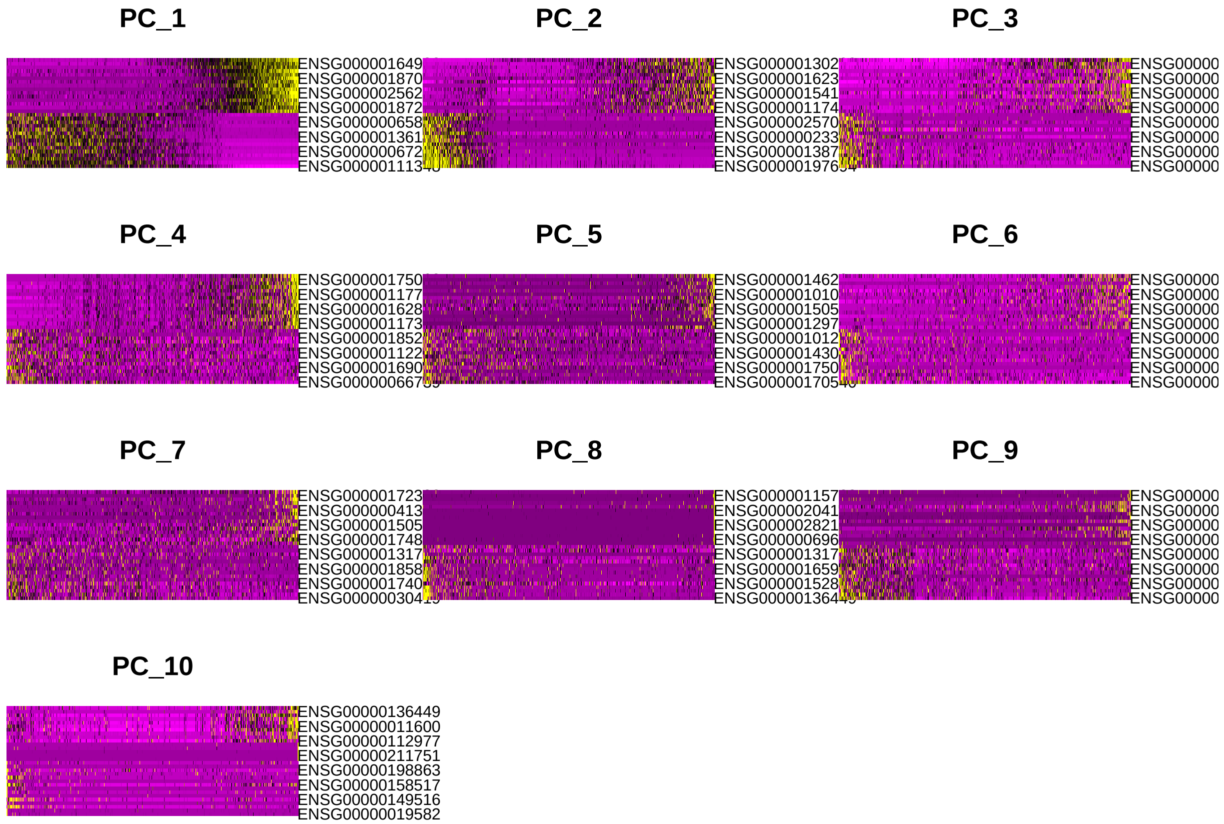


Cell Cycle Scoring uses default Seurat G2M and S genes of interest to assign each cell a score, based on its expression of G2/M and S phase markers. Cells expressing neither are likely not cycling and in G1 phase as the data set should be anticorrelated.

```
rnaseq_data_score_seuratgenes <- CellCycleScoring(rnaseq_data_var_seuratgenes, s.features = seurat.s.ge
```

```
## Warning: The following features are not present in the object: ENSG00000112312,
## not searching for symbol synonyms
```

```
## Warning: The following features are not present in the object: ENSG00000129195,
## ENSG00000189159, not searching for symbol synonyms
```

A PCA is generated from scored data without dimensionality reduction

```
DimPlot(object = rnaseq_data_score_seuratgenes, combine = FALSE, cols = c("red", "steelblue", "darkgree
```

```
## [[1]]
```

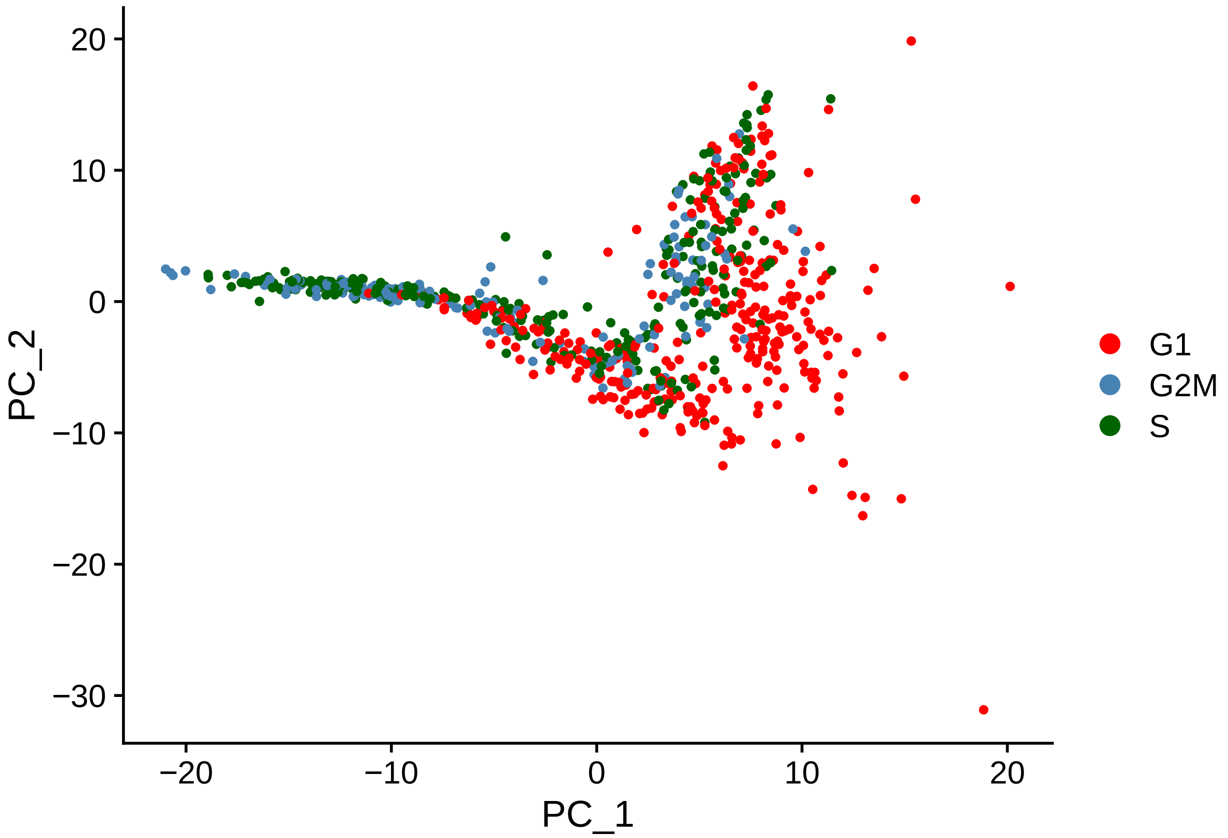


The top six results are printed to ensure proper output of phase assignment, S.Score and G2M.Score

```
head(rnaseq_data_score_seuratgenes[[]])
```

```
##          orig.ident nCount_RNA nFeature_RNA     S.Score G2M.Score Phase
## Prog_013       Prog    2029157         8366 -0.50854828 -1.381708    G1
## Prog_019       Prog    2408705         8194 -0.90542682  1.477430   G2M
## Prog_031       Prog    1033611         8349 -1.77053514 -1.295574    G1
## Prog_037       Prog    1068895         7955 -1.50509622  2.350027   G2M
## Prog_008       Prog    3174838         8596  2.37167494  0.304161     S
## Prog_014       Prog    1971269         8957 -0.07372224  1.050870   G2M
##          old.ident
## Prog_013      Prog
## Prog_019      Prog
## Prog_031      Prog
## Prog_037      Prog
## Prog_008      Prog
## Prog_014      Prog
```

Ridge plots of commonly seen genes are generated to ensure the proper differentiation of high S and high G2M scoring genes

```
RidgePlot(rnaseq_data_score_seuratgenes, features = c("ENSG00000117399", "ENSG00000169679", "ENSG0000001
```

```
## Picking joint bandwidth of 0.5
```

```
## Picking joint bandwidth of 0.216

## Picking joint bandwidth of 0.59

## Picking joint bandwidth of 0.0478
```

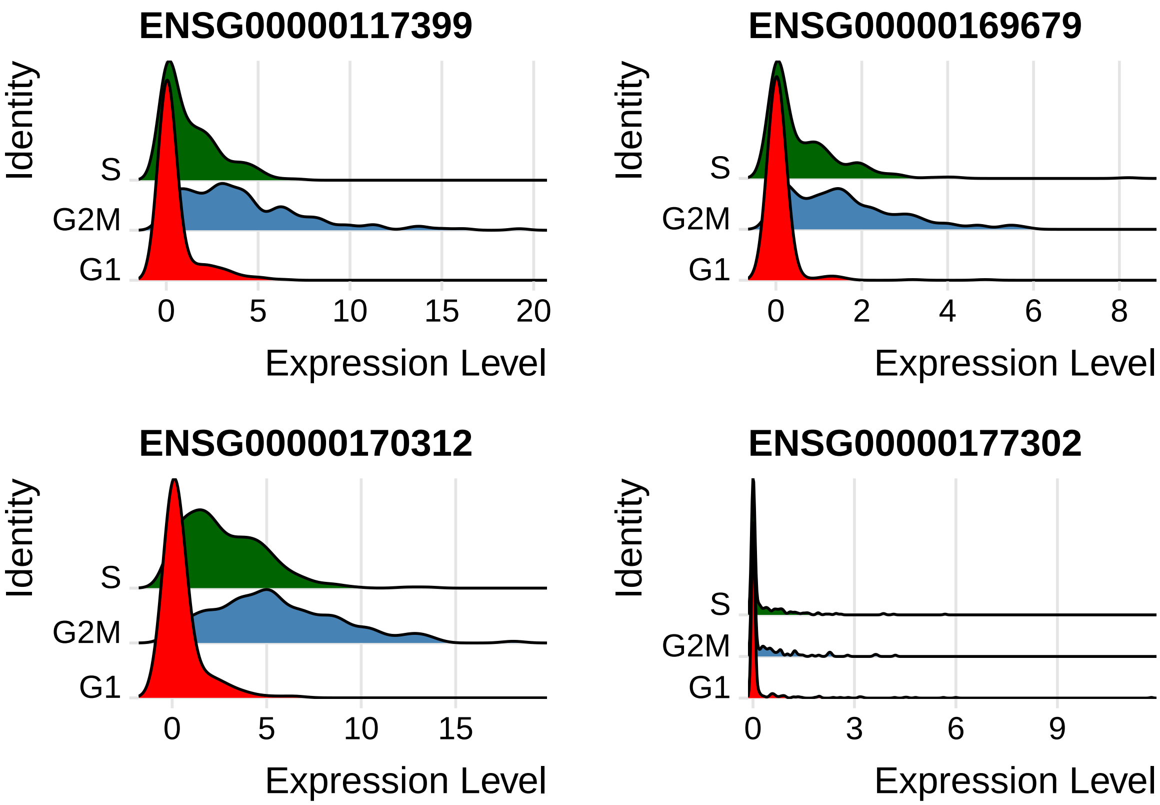


Generated and plotted PCA with proper grouping of phase scoring are produced

```
rnaseq_data_score_seuratgenes_pca <- RunPCA(rnaseq_data_score_seuratgenes, npcs = (npcs_use), features =
```

```
## Warning in PrepDR(object = object, features = features, verbose = verbose): The
## following 3 features requested have not been scaled (running reduction without
## them): ENSG00000112312, ENSG00000129195, ENSG00000189159

## Warning in irlba(A = t(x = object), nv = npcs, ...): You're computing too large
## a percentage of total singular values, use a standard svd instead.

## PC_ 1
## Positive:  ENSG00000089685, ENSG00000111665, ENSG00000148773, ENSG00000170312, ENSG00000131747, ENSG
##     ENSG00000137804, ENSG00000013810, ENSG00000138160, ENSG00000113810, ENSG00000178999, ENSG0000008
##     ENSG00000167325, ENSG00000169679, ENSG00000142945, ENSG00000117724, ENSG00000117650, ENSG0000012
## Negative:  ENSG00000158402, ENSG00000174371, ENSG00000144354, ENSG00000076248, ENSG00000169607, ENSG
##     ENSG00000010292, ENSG00000118412, ENSG00000123975, ENSG00000134222, ENSG00000129173, ENSG0000012
##     ENSG00000102974, ENSG00000076003, ENSG00000119969, ENSG00000159259, ENSG00000092853, ENSG0000010
```

```
## PC_ 2
## Positive:  ENSG00000175063, ENSG00000137804, ENSG00000117724, ENSG00000142945, ENSG00000088325, ENSG[illegible]
##     ENSG00000087586, ENSG00000134690, ENSG00000115163, ENSG00000092140, ENSG00000138182, ENSG0000014[illegible]
##     ENSG00000131747, ENSG00000158402, ENSG00000136108, ENSG00000111665, ENSG00000011426, ENSG0000010[illegible]
## Negative:  ENSG00000132646, ENSG00000276043, ENSG00000100297, ENSG00000076003, ENSG00000075131, ENSG[illegible]
##     ENSG00000094804, ENSG00000131153, ENSG00000175305, ENSG00000119969, ENSG00000076248, ENSG0000016[illegible]
##     ENSG00000168496, ENSG00000176890, ENSG00000159259, ENSG00000143476, ENSG00000111247, ENSG0000015[illegible]
## PC_ 3
## Positive:  ENSG00000131747, ENSG00000101868, ENSG00000049541, ENSG00000178999, ENSG00000176890, ENSG[illegible]
##     ENSG00000148773, ENSG00000094916, ENSG00000092470, ENSG00000102974, ENSG00000011426, ENSG0000011[illegible]
##     ENSG00000170312, ENSG00000175216, ENSG00000051180, ENSG00000137807, ENSG00000143476, ENSG0000012[illegible]
## Negative:  ENSG00000115163, ENSG00000157456, ENSG00000094804, ENSG00000175305, ENSG00000168496, ENSG[illegible]
##     ENSG00000092853, ENSG00000139354, ENSG00000143815, ENSG00000112742, ENSG00000100401, ENSG0000011[illegible]
##     ENSG00000075131, ENSG00000142945, ENSG00000089685, ENSG00000072571, ENSG00000010292, ENSG0000008[illegible]
## PC_ 4
## Positive:  ENSG00000076248, ENSG00000143476, ENSG00000119969, ENSG00000049541, ENSG00000136108, ENSG[illegible]
##     ENSG00000137807, ENSG00000117650, ENSG00000100297, ENSG00000104738, ENSG00000010292, ENSG0000014[illegible]
##     ENSG00000137804, ENSG00000143815, ENSG00000175063, ENSG00000088325, ENSG00000114346, ENSG0000012[illegible]
## Negative:  ENSG00000112742, ENSG00000176890, ENSG00000178999, ENSG00000089685, ENSG00000111665, ENSG[illegible]
##     ENSG00000143228, ENSG00000171848, ENSG00000188229, ENSG00000148773, ENSG00000132646, ENSG0000016[illegible]
##     ENSG00000102974, ENSG00000092853, ENSG00000197299, ENSG00000169607, ENSG00000101868, ENSG0000015[illegible]
## PC_ 5
## Positive:  ENSG00000143815, ENSG00000095002, ENSG00000159259, ENSG00000168496, ENSG00000077514, ENSG[illegible]
##     ENSG00000049541, ENSG00000137804, ENSG00000166508, ENSG00000013810, ENSG00000137807, ENSG0000011[illegible]
##     ENSG00000163950, ENSG00000171848, ENSG00000158402, ENSG00000170312, ENSG00000089685, ENSG0000011[illegible]
## Negative:  ENSG00000171421, ENSG00000162607, ENSG00000125630, ENSG00000143401, ENSG00000197299, ENSG[illegible]
##     ENSG00000173207, ENSG00000132780, ENSG00000092853, ENSG00000115163, ENSG00000120802, ENSG0000016[illegible]
##     ENSG00000118412, ENSG00000113810, ENSG00000112742, ENSG00000175216, ENSG00000157456, ENSG0000014[illegible]
```

```r
DimPlot(rnaseq_data_score_seuratgenes_pca,combine = FALSE, cols = c("red", "steelblue", "darkgreen"))
```

```
## [[1]]
```

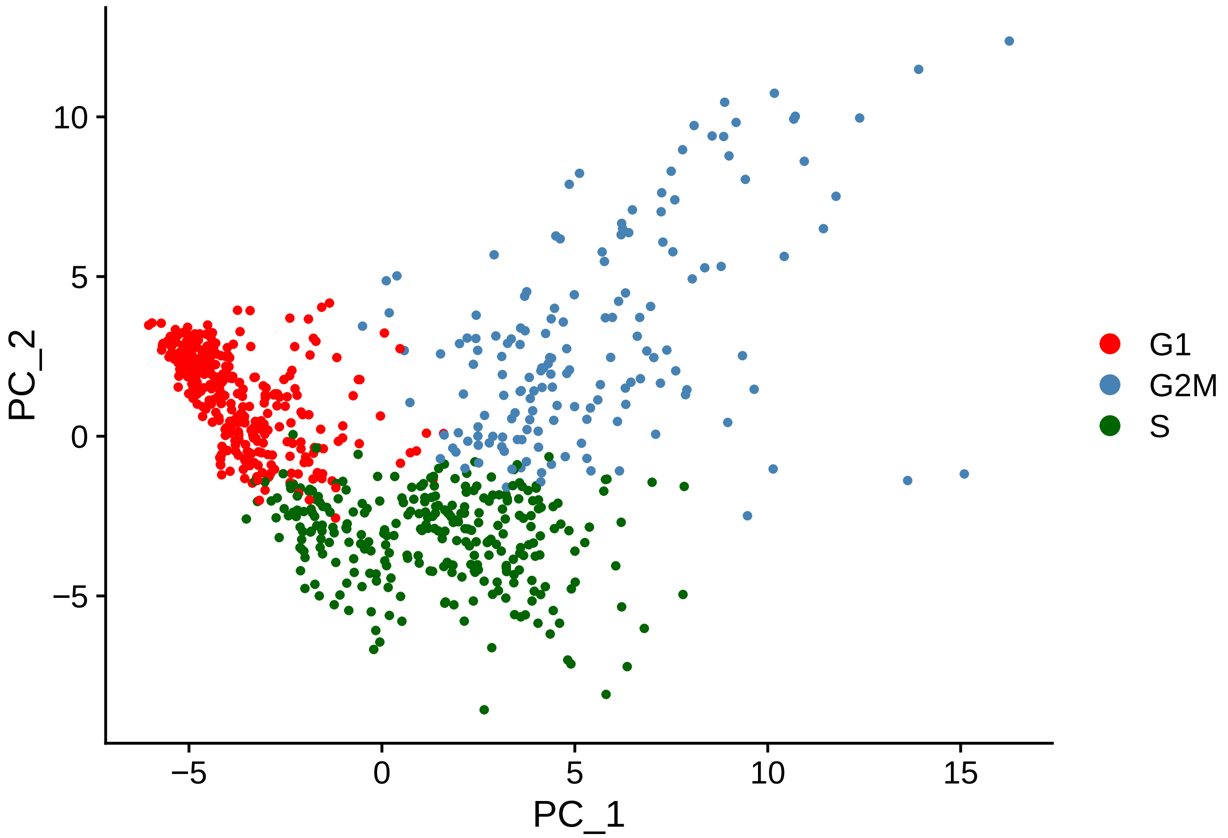


The phase percentages of G1, S and G2M are calculated and plotted

```
#Calculates and plots values of phase percentages of G1, S and G2M
phase_df <- rnaseq_data_score_seuratgenes_pca$Phase
phase_df <- count(phase_df)
names(phase_df)[1] <- 'Phase'
sumphase <- sum(phase_df$freq)
percentage_df <- phase_df$freq /sumphase *100
phase_df$Percentage <- percentage_df
phase_df <- phase_df %>%
  arrange(desc(Phase)) %>%
  mutate(prop = freq / sum(phase_df$freq) *100) %>%
  mutate(ypos = cumsum(prop)- 0.5*prop )
percent=(phase_df$Percentage)
percent=round(percent,digits=3)
phase_df$PercentageLabel <- paste0((percent), "%")

#Plots as a pie chart the assigned phase percentages in the overall intial count matrix
ggplot(data = phase_df, aes(x = "", y = Percentage, fill = Phase)) +
  geom_bar(stat = "identity") +
  coord_polar("y")  + geom_text(aes(y = ypos, label = PercentageLabel), color = "white", size=3)+ scale_
```

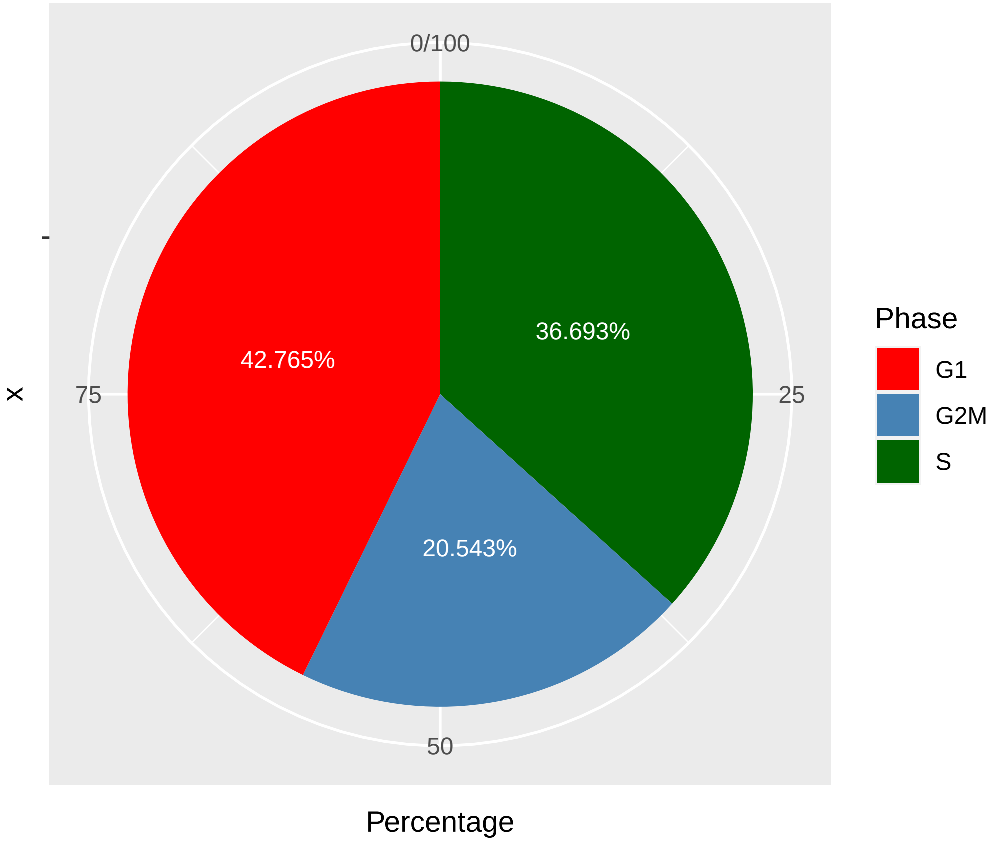


#### Tidying up data for G2 and M Seurat run We then isolate and subsets G2M, S and G1 into seperate variables so G2M can be processed further

```
rnaseq_seuratgenes_PCA_PhaseData <- FetchData(object = rnaseq_data_score_seuratgenes, vars = c('orig.id
rnaseq_seuratgenes_PCA_PhaseData<-tibble::rownames_to_column(rnaseq_seuratgenes_PCA_PhaseData, "run")
rnaseq_seuratgenes_G2M <- filter(rnaseq_seuratgenes_PCA_PhaseData, Phase == "G2M")
rnaseq_seuratgenes_S <-  filter(rnaseq_seuratgenes_PCA_PhaseData, Phase == "S")
rnaseq_seuratgenes_G1 <-  filter(rnaseq_seuratgenes_PCA_PhaseData, Phase == "G1")
```

The G2M scored data is tidied up which removes extra data so the G2M only fraction of input data can be ran through a modified Seurat phase assignment chunk to separate out a mitotic specific fraction

```
read_data_rotate <- data.frame(t(read_data[]))
read_data_rotate <- tibble::rownames_to_column(read_data_rotate)
names(read_data_rotate)[names(read_data_rotate) == "rowname"] <- "run"

joinfilter_g2m <- tibble::rownames_to_column(rnaseq_seuratgenes_G2M)

g2m_filtercells <- joinfilter_g2m$run
g2m_readdata <- filter(read_data_rotate, run %in% g2m_filtercells)
g2m_readdata <- data.frame(t(g2m_readdata[]))
names(g2m_readdata) <- as.matrix(g2m_readdata[1, ])
g2m_readdata <- g2m_readdata[-1, ]
```

#### Modification to CellCycleScoring for use in G2 and M phase assignment

Here we show the Seurat Cell Cycle Scoring function with the ability to assign G1 phase cells removed as this function would interfere with assigning phase to the G2/M subsetted cells into G2 and M specifically and ensure the proper labelling of said cells

```
CellCycleScoring_G1Disable <- function(
  object,
  s.features,
  g2m.features,
  ctrl = NULL,
  set.ident = FALSE,
  ...
) {
  name <- 'Cell.Cycle'
  features <- list('S.Score' = s.features, 'G2M.Score' = g2m.features)
  if (is.null(x = ctrl)) {
    ctrl <- min(vapply(X = features, FUN = length, FUN.VALUE = numeric(length = 1)))
  }
  object.cc <- AddModuleScore(
    object = object,
    features = features,
    name = name,
    ctrl = ctrl,
    ...
  )
  cc.columns <- grep(pattern = name, x = colnames(x = object.cc[[]]), value = TRUE)
  cc.scores <- object.cc[[cc.columns]]
  rm(object.cc)
  CheckGC()
  assignments <- apply(
    X = cc.scores,
    MARGIN = 1,
    FUN = function(scores, first = 'G2', second = 'M', null = 'G1') {
        if (length(which(x = scores == max(scores))) > 1) {
          return('Undecided')
        } else {
          return(c(first, second)[which(x = scores == max(scores))])
        }
      }
  )
  cc.scores <- merge(x = cc.scores, y = data.frame(assignments), by = 0)
  colnames(x = cc.scores) <- c('rownames', 'S.Score', 'G2M.Score', 'Phase')
  rownames(x = cc.scores) <- cc.scores$rownames
  cc.scores <- cc.scores[, c('S.Score', 'G2M.Score', 'Phase')]
  object[[colnames(x = cc.scores)]] <- cc.scores
  if (set.ident) {
    object[['old.ident']] <- Idents(object = object)
    Idents(object = object) <- 'Phase'
  }
  return(object)
}
```

#### G2 and M Seurat phase assignment

A second Seurat cell cycle phase assignment is run on log normalized cells using the generated Interphase and Mitotic gene list of interest, derived from the differentially expressed genes tested via Bulk RNA sequencing

```
#Reassigns column numbers for npcs used in RunPCA Functions for the G2M only subsetted fraction
npcs_num <- ncol(g2m_readdata)

if (npcs_num  < 50) {
npcs_use_g2m <- npcs_num-1
} else {
npcs_use_g2m <- 50
}
```

Chosen input in this step uses generated gene list so the points of variance for phase assignment are Interphase (Regarded as G2 as they are derived from a G2M only fraction) and M Phase fraction

```
cell_marker_generated <- read_csv("data/RNAseq_sigGene_MvsI_Ensembl.csv")
```

```
## Rows: 27 Columns: 2
## -- Column specification --------------------------------------------------------
## Delimiter: ","
## chr (2): Interphase_Padj, Mitotic_Padj
##
## i Use `spec()` to retrieve the full column specification for this data.
## i Specify the column types or set `show_col_types = FALSE` to quiet this message.
```

```
g2_genes <- cell_marker_generated$Interphase_Padj
m_genes <- cell_marker_generated$Mitotic_Padj
g2_genes <- na.omit(g2_genes)
m_genes <- na.omit(m_genes)
```

A Seurat object is created from raw data

```
#Create a Seurat object from raw data
rnaseq_data_generatedgenes <- CreateSeuratObject(counts = g2m_readdata)
```

Normalisation step of count matrix using Relative counts, input G2 and M feature counts for each cell are divided bt total counts and scaled by scale.factor. This is then natural-log transformed using log1p to now accurately scale the G2 and M variables respectively.

```
rnaseq_data_var_generatedgenes <- NormalizeData(
  rnaseq_data_generatedgenes,
  assay = "RNA",
  normalization.method = "LogNormalize",
  scale.factor = 10000,
  margin = 1,
  verbose = TRUE
)
```

A mean variability plot is used to assign outliers in data set, the selection method vst was chosen. Vst fits the relationship of log(variance) and log(mean) using local polynomial regression (loess). The function then standardizes the feature valued using the mean and expected variance from that fitted relationship. The standardized values are used to calculate the feature variance. In this step as stated G2 and M rather than S and G2M.

```
rnaseq_data_var_generatedgenes <- FindVariableFeatures(rnaseq_data_var_generatedgenes, selection.method
```

Features are centered and scaled in the dataset.

```
rnaseq_data_var_generatedgenes <- ScaleData(rnaseq_data_var_generatedgenes, features = rownames(rnaseq_
```

```
## Centering and scaling data matrix
```

Post scaling a PCA dimensionality reduction is run. Uses npcs assigned at chunk start and uses variable features of input count matrix with default seurat gene list of interest. PrintPCAParams can be run for more detail.

```
rnaseq_data_var_generatedgenes <- RunPCA(rnaseq_data_var_generatedgenes, features = VariableFeatures(rn
```

```
## PC_ 1
## Positive:  ENSG00000174804, ENSG00000142669, ENSG00000089327, ENSG00000145287, ENSG00000153551, ENSG
## Negative:  ENSG00000167741, ENSG00000164932, ENSG00000164010, ENSG00000107789, ENSG00000187010, ENSG
## PC_ 2
## Positive:  ENSG00000130203, ENSG00000179348, ENSG00000011295, ENSG00000186766, ENSG00000005961, ENSG
## Negative:  ENSG00000124216, ENSG00000112799, ENSG00000110375, ENSG00000101544, ENSG00000140471, ENSG
## PC_ 3
## Positive:  ENSG00000168685, ENSG00000113312, ENSG00000143344, ENSG00000185437, ENSG00000182866, ENSG
## Negative:  ENSG00000197561, ENSG00000085491, ENSG00000140749, ENSG00000257017, ENSG00000197694, ENSG
## PC_ 4
## Positive:  ENSG00000141076, ENSG00000134061, ENSG00000091157, ENSG00000108861, ENSG00000112576, ENSG
## Negative:  ENSG00000243063, ENSG00000169684, ENSG00000100147, ENSG00000172426, ENSG00000168878, ENSG
## PC_ 5
## Positive:  ENSG00000006638, ENSG00000197943, ENSG00000135253, ENSG00000183615, ENSG00000108861, ENSG
## Negative:  ENSG00000107036, ENSG00000262406, ENSG00000198771, ENSG00000142619, ENSG00000148814, ENSG
## PC_ 6
## Positive:  ENSG00000146281, ENSG00000163737, ENSG00000172382, ENSG00000166831, ENSG00000120457, ENSG
## Negative:  ENSG00000137845, ENSG00000117399, ENSG00000224189, ENSG00000094975, ENSG00000163751, ENSG
## PC_ 7
## Positive:  ENSG00000215612, ENSG00000106077, ENSG00000076928, ENSG00000090339, ENSG00000273749, ENSG
## Negative:  ENSG00000102760, ENSG00000236279, ENSG00000125730, ENSG00000063660, ENSG00000149516, ENSG
## PC_ 8
## Positive:  ENSG00000119431, ENSG00000256590, ENSG00000072840, ENSG00000170476, ENSG00000175063, ENSG
## Negative:  ENSG00000113312, ENSG00000179673, ENSG00000228741, ENSG00000182957, ENSG00000273167, ENSG
## PC_ 9
## Positive:  ENSG00000138161, ENSG00000167900, ENSG00000001497, ENSG00000067836, ENSG00000219607, ENSG
## Negative:  ENSG00000124191, ENSG00000125730, ENSG00000144677, ENSG00000106066, ENSG00000128944, ENSG
## PC_ 10
## Positive:  ENSG00000176571, ENSG00000147168, ENSG00000104341, ENSG00000123243, ENSG00000147852, ENSG
## Negative:  ENSG00000124571, ENSG00000048462, ENSG00000256590, ENSG00000119431, ENSG00000170476, ENSG
```

A heat map of the points of variance PC1 to PC10 are plotted

```r
DimHeatmap(rnaseq_data_var_generatedgenes, dims = c(1:10))
```

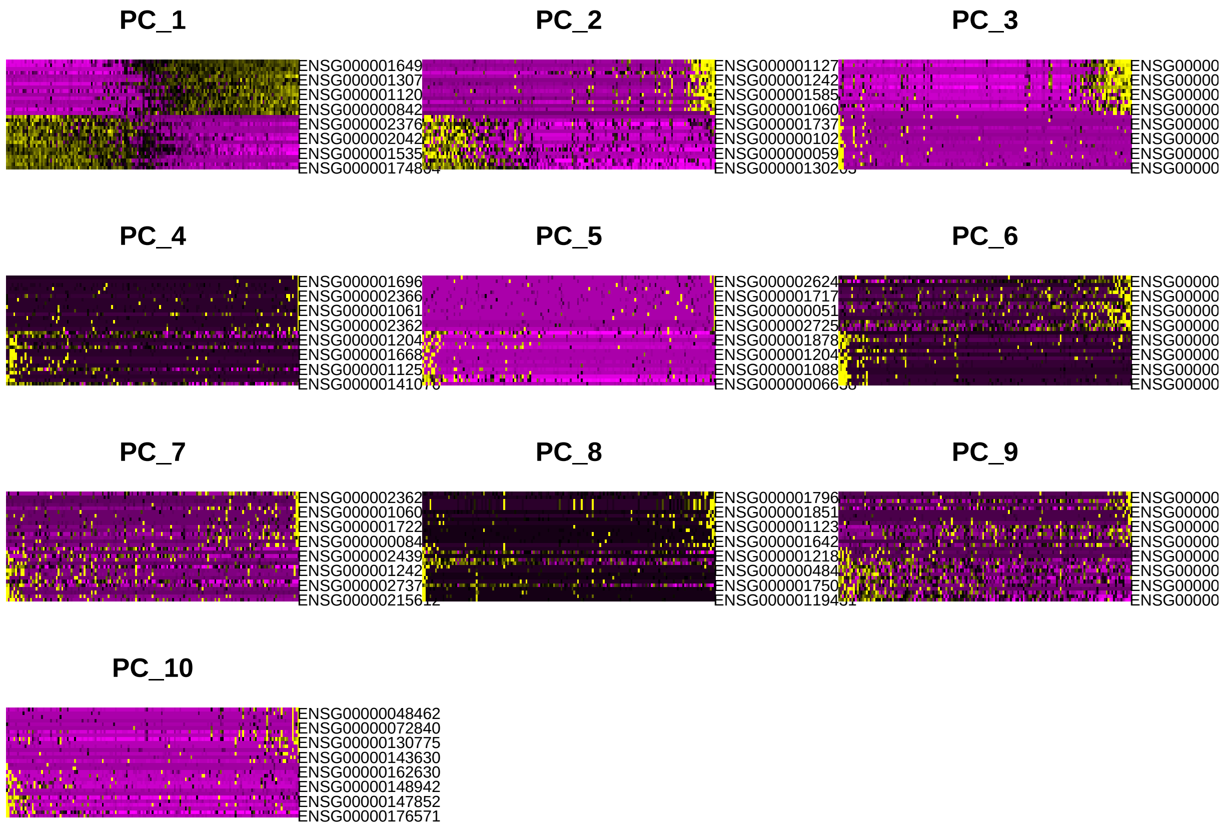


Cell Cycle Scoring uses CellCycleScoring_G1Disable to correctly assign cells based on its expression of G2 (Interphase) and M phase markers, G1 calling is not required in this step.

```r
rnaseq_data_score_generatedgenes <- CellCycleScoring_G1Disable(rnaseq_data_var_generatedgenes, s.featur
```

```
## Warning: The following features are not present in the object: ENSG00000273759,
## ENSG00000165244, ENSG00000272106, ENSG00000275484, ENSG00000165948,
## ENSG00000207547, not searching for symbol synonyms
```

```
## Warning: The following features are not present in the object: ENSG00000129195,
## ENSG00000163535, ENSG00000024526, ENSG00000090889, not searching for symbol
## synonyms
```

A PCA is generated from scored data without dimensionality reduction

```r
DimPlot(object = rnaseq_data_score_generatedgenes, combine = FALSE, cols = c("#E3E857", "#6a329f", "#8f
```

```
## [[1]]
```

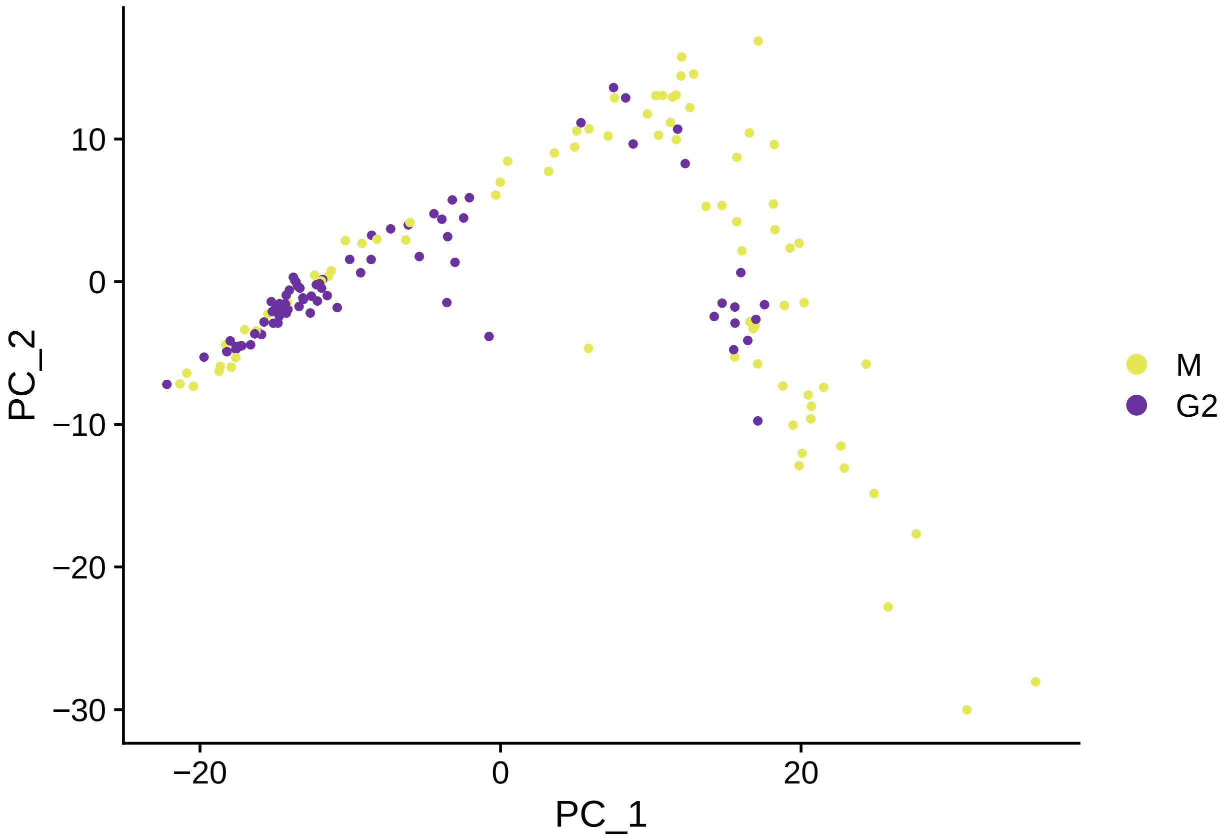


The top six results are printed to ensure proper output of phase assignment, S.Score (Modified to investigate G2 genes of interest) and G2M.Score (Modified to investigate M genes of interest)

```
head(rnaseq_data_score_generatedgenes[[]])
```

```
##          orig.ident nCount_RNA nFeature_RNA     S.Score   G2M.Score Phase
## Prog_019       Prog    2408705         8194 -0.17581863  0.17891918     M
## Prog_037       Prog    1068895         7955 -0.38620260  0.35113848     M
## Prog_014       Prog    1971269         8957  0.02925160 -0.07011165    G2
## Prog_032       Prog    1260663         8301  0.08545737 -0.16467122    G2
## Prog_038       Prog    1754706         8132  0.06324387 -0.29006825    G2
## Prog_009       Prog    3551293         8694  0.09625127 -0.34645865    G2
##          old.ident
## Prog_019      Prog
## Prog_037      Prog
## Prog_014      Prog
## Prog_032      Prog
## Prog_038      Prog
## Prog_009      Prog
```

Generated and plotted PCA with proper grouping of phase scoring are produced

```
rnaseq_data_score_generatedgenes_pca <- RunPCA(rnaseq_data_score_generatedgenes, npcs = (npcs_use), fea
```

```
## Warning in PrepDR(object = object, features = features, verbose = verbose): The
```

```
## following 10 features requested have not been scaled (running reduction without
## them): ENSG00000273759, ENSG00000165244, ENSG00000272106, ENSG00000275484,
## ENSG00000165948, ENSG00000207547, ENSG00000129195, ENSG00000163535,
## ENSG00000024526, ENSG00000090889

## Warning in irlba(A = t(x = object), nv = npcs, ...): You're computing too large
## a percentage of total singular values, use a standard svd instead.

## Warning in irlba(A = t(x = object), nv = npcs, ...): did not converge--results
## might be invalid!; try increasing work or maxit

## Warning: Requested number is larger than the number of available items (35).
## Setting to 35.

## Warning: Requested number is larger than the number of available items (35).
## Setting to 35.

## Warning: Requested number is larger than the number of available items (35).
## Setting to 35.

## Warning: Requested number is larger than the number of available items (35).
## Setting to 35.

## Warning: Requested number is larger than the number of available items (35).
## Setting to 35.

## PC_ 1
## Positive:  ENSG00000105173, ENSG00000165879, ENSG00000094804, ENSG00000076248, ENSG00000101412, ENSG[illegible]
##     ENSG00000175643, ENSG00000173218, ENSG00000119938, ENSG00000138778, ENSG00000118193, ENSG000001[illegible]
## Negative:  ENSG00000170540, ENSG00000137804, ENSG00000128944, ENSG00000117399, ENSG00000088325, ENSG[illegible]
##     ENSG00000126787, ENSG00000068489, ENSG00000117650, ENSG00000166851, ENSG00000112984, ENSG0000014[illegible]
## PC_ 2
## Positive:  ENSG00000105173, ENSG00000166851, ENSG00000094804, ENSG00000145386, ENSG00000100297, ENSG[illegible]
##     ENSG00000118193, ENSG00000165879, ENSG00000066279, ENSG00000112984, ENSG00000137812, ENSG0000007[illegible]
## Negative:  ENSG00000138180, ENSG00000175643, ENSG00000138778, ENSG00000108306, ENSG00000068489, ENSG[illegible]
##     ENSG00000143476, ENSG00000143228, ENSG00000137804, ENSG00000170540, ENSG00000088325, ENSG0000016[illegible]
## PC_ 3
## Positive:  ENSG00000137812, ENSG00000118193, ENSG00000108306, ENSG00000119938, ENSG00000139354, ENSG[illegible]
##     ENSG00000138182, ENSG00000066279, ENSG00000166851, ENSG00000169679, ENSG00000138778, ENSG0000009[illegible]
## Negative:  ENSG00000143476, ENSG00000175643, ENSG00000100297, ENSG00000117650, ENSG00000101412, ENSG[illegible]
##     ENSG00000143228, ENSG00000137804, ENSG00000173218, ENSG00000145386, ENSG00000117399, ENSG0000007[illegible]
## PC_ 4
## Positive:  ENSG00000173218, ENSG00000066279, ENSG00000119938, ENSG00000175643, ENSG00000138778, ENSG[illegible]
##     ENSG00000170540, ENSG00000186193, ENSG00000101412, ENSG00000072571, ENSG00000108306, ENSG0000013[illegible]
## Negative:  ENSG00000145386, ENSG00000117399, ENSG00000138180, ENSG00000126787, ENSG00000128944, ENSG[illegible]
##     ENSG00000143228, ENSG00000117650, ENSG00000204256, ENSG00000118193, ENSG00000100297, ENSG0000016[illegible]
## PC_ 5
## Positive:  ENSG00000143228, ENSG00000068489, ENSG00000204256, ENSG00000139354, ENSG00000175643, ENSG[illegible]
##     ENSG00000126787, ENSG00000105173, ENSG00000138180, ENSG00000170540, ENSG00000143476, ENSG0000013[illegible]
## Negative:  ENSG00000101412, ENSG00000186193, ENSG00000128944, ENSG00000173218, ENSG00000117650, ENSG[illegible]
##     ENSG00000108306, ENSG00000072571, ENSG00000138182, ENSG00000165879, ENSG00000165724, ENSG0000011[illegible]
```

```r
DimPlot(object = rnaseq_data_score_generatedgenes_pca, combine = FALSE, cols = c("#984DD3", "#E88A13",
```

```
## [[1]]
```

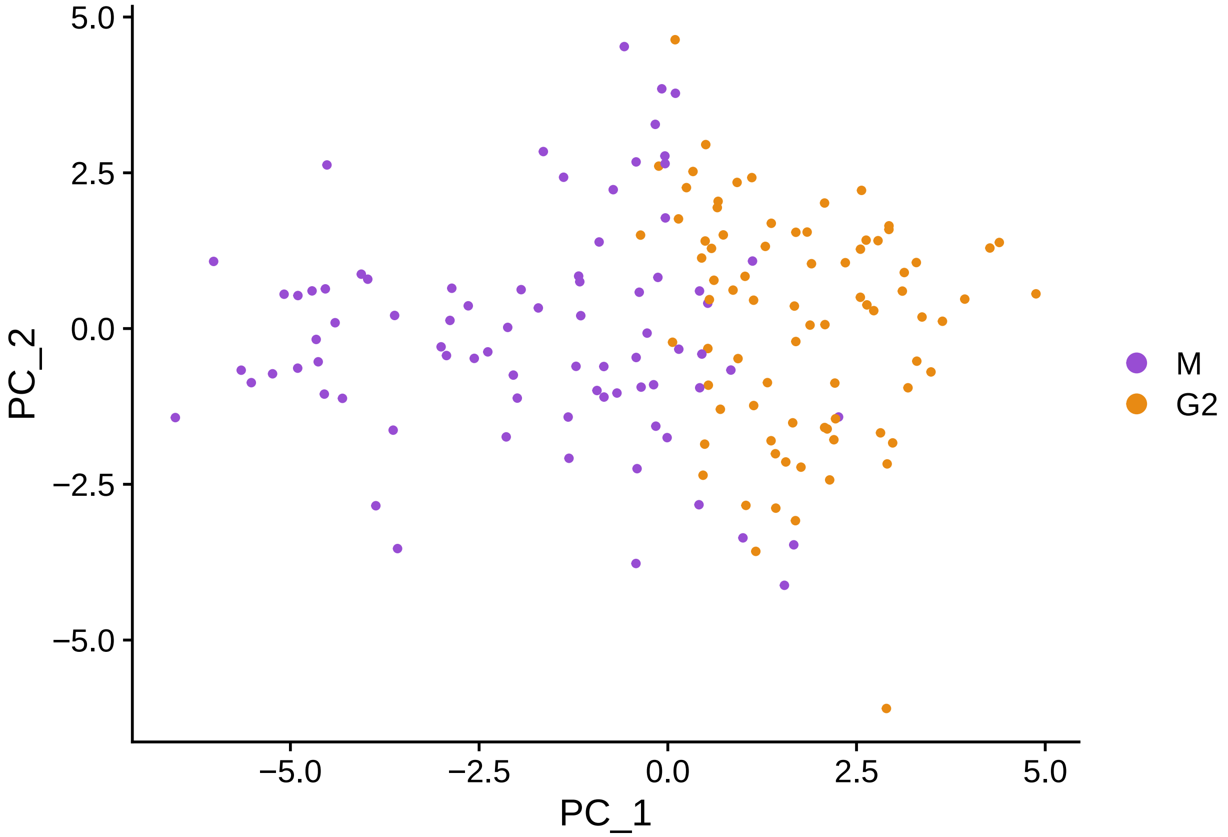


Plots commonly seen genes to ensure the proper expression of high G2 and high M scoring genes is present

```r
#Plots commonly seen genes to ensure the proper differentiation of high G2 (representing the G2 cells)
RidgePlot(rnaseq_data_score_generatedgenes, features = c("ENSG00000117399", "ENSG00000169679", "ENSG000
```

```
## Picking joint bandwidth of 0.228

## Picking joint bandwidth of 0.19

## Picking joint bandwidth of 0.193

## Picking joint bandwidth of 0.0714
```

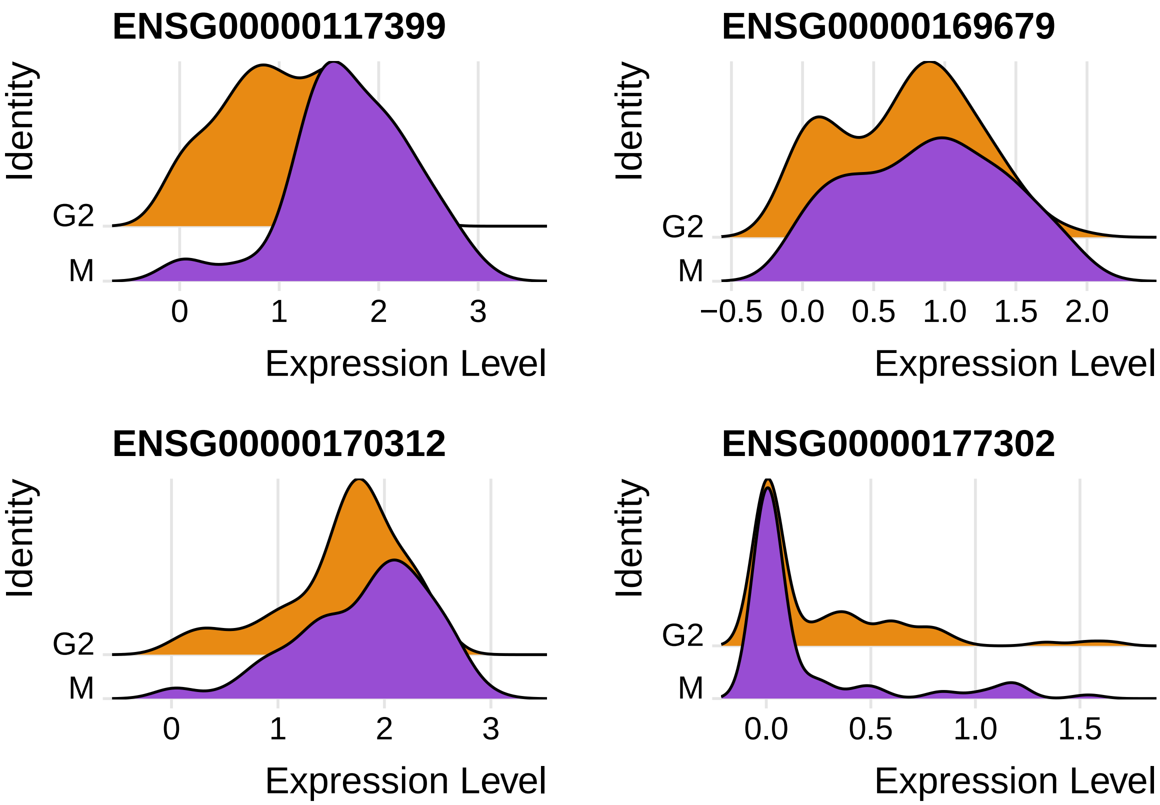


The phase percentages of G2 and M are calculated and plotted

```
phase_geninput <- rnaseq_data_score_generatedgenes$Phase
phase_geninput <- count(phase_geninput)
names(phase_geninput)[1] <- 'Phase'
sumphase <- sum(phase_geninput$freq)
percentage_df <- phase_geninput$freq /sumphase *100
phase_geninput$Percentage <- percentage_df
phase_geninput <- phase_geninput %>%
  arrange(desc(Phase)) %>%
  mutate(prop = freq / sum(phase_geninput$freq) *100) %>%
  mutate(ypos = cumsum(prop)- 0.5*prop )
percent=(phase_geninput$Percentage)
percent=round(percent,digits=3)
phase_geninput$PercentageLabel <- paste0((percent), "%")

#Plots as a pie chart the assigned phase percentages in the overall intial count matrix
ggplot(data = phase_geninput, aes(x = "", y = Percentage, fill = Phase)) +
  geom_bar(stat = "identity") +
  coord_polar("y")  + geom_text(aes(y = ypos, label = PercentageLabel), color = "white", size=3) + scal
```

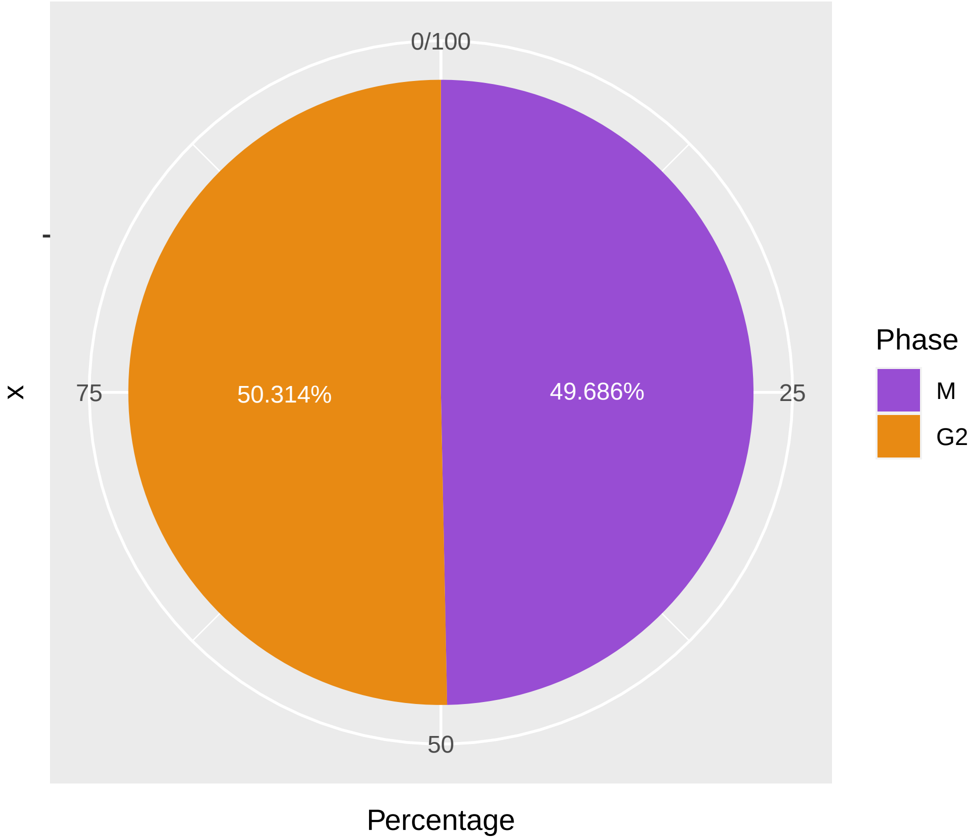


To ensure that there is no error in naming we added this function to ensure that the G2 and M assigned cells in the Seurat G2 and M phase assignment would always be correct

```
g2m_cellcyclescored <- FetchData(object = rnaseq_data_score_generatedgenes, vars = c('orig.ident', 'nCou
g2m_cellcyclescored$Phase <- revalue(g2m_cellcyclescored$Phase, c("G2M"="M"))
```

```
## The following `from` values were not present in `x`: G2M
```

```
g2m_cellcyclescored$Phase <- revalue(g2m_cellcyclescored$Phase, c("S"="G2"))
```

```
## The following `from` values were not present in `x`: S
```

The G2 and M scored cells are the rejoined with the intial count matrix to generated a new matrix containing only G2 and M scored genes post phase assignment from the G2 and M Seurat sort

```
#The now renamed G2 and M scored cells are the rejoined with the intial count matrix to generated a new
g2_m_finalsplit <- g2m_cellcyclescored %>% rownames_to_column("run")
g2_m_phase_Counts <- left_join(g2_m_finalsplit, read_data_rotate, by = "run")
g2_m_phase_Counts <- subset(g2_m_phase_Counts, select = -c(orig.ident, nCount_RNA, nFeature_RNA, S.Scor
g2_m_phase_Counts <- data.frame(t(g2_m_phase_Counts[]))
names(g2_m_phase_Counts) <- as.matrix(g2_m_phase_Counts[1, ])
g2_m_phase_Counts <- g2_m_phase_Counts[-1, ]
```

If required this write out function can be enabled to generate this G2 and M specific count matrix

```
write.csv(g2_m_phase_Counts, file = (paste((GSE),"G2_and_M_Phases_Assigned.csv")))
```

##### Collation of phase assignment and graphing

Here we combine all phase assigned variables together to get a count matrix with G1, G2, S and M assigned cells and write out a complete count matrix with phase assignment attached

```
#Combined all phase assigned variables together to get a count matrix with G1, G2, S and M assigned cel
phasescored_allruns <- rbind(rnaseq_seuratgenes_S,rnaseq_seuratgenes_G1,g2_m_finalsplit)
phasescored_allruns_countmatrix <- left_join(phasescored_allruns, read_data_rotate, by = "run")
phasescored_allruns_countmatrix <- subset(phasescored_allruns_countmatrix, select = -c (orig.ident, nCou
phasescored_allruns_countmatrix <- data.frame(t(phasescored_allruns_countmatrix[]))
names(phasescored_allruns_countmatrix) <- as.matrix(phasescored_allruns_countmatrix[1, ])
phasescored_allruns_countmatrix <- phasescored_allruns_countmatrix[-1, ]

write.csv(phasescored_allruns_countmatrix, file = (paste((GSE),"All_Phases_Assigned.csv")))
```

Finally we calculate the assigned phase percentages in the complete G1, G2, S and M assigned count matrix and graph these percentages

```
#Calculates the assigned phase percentages in the overall intial count matrix
phasescored_allruns_countmatrix <- data.frame(t(phasescored_allruns_countmatrix[]))
phase_df1 <- phasescored_allruns_countmatrix$Phase
phase_df1 <- count(phase_df1)
names(phase_df1)[1] <- 'Phase'
sumphase <- sum(phase_df1$freq)
percentage_df <- phase_df1$freq /sumphase *100
phase_df1$Percentage <- percentage_df
phase_df1 <- phase_df1 %>%
  arrange(desc(Phase)) %>%
  mutate(prop = freq / sum(phase_df1$freq) *100) %>%
  mutate(ypos = cumsum(prop)- 0.5*prop )
percent=(phase_df1$Percentage)
percent=round(percent,digits=3)
phase_df1$PercentageLabel <- paste0((percent), "%")


#Plots as a pie chart the assigned phase percentages in the overall intial count matrix
ggplot(data = phase_df1, aes(x = "", y = Percentage, fill = Phase)) +
  geom_bar(stat = "identity") +
  coord_polar("y")  + geom_text(aes(y = ypos, label = PercentageLabel), color = "white", size=3) + scal
```

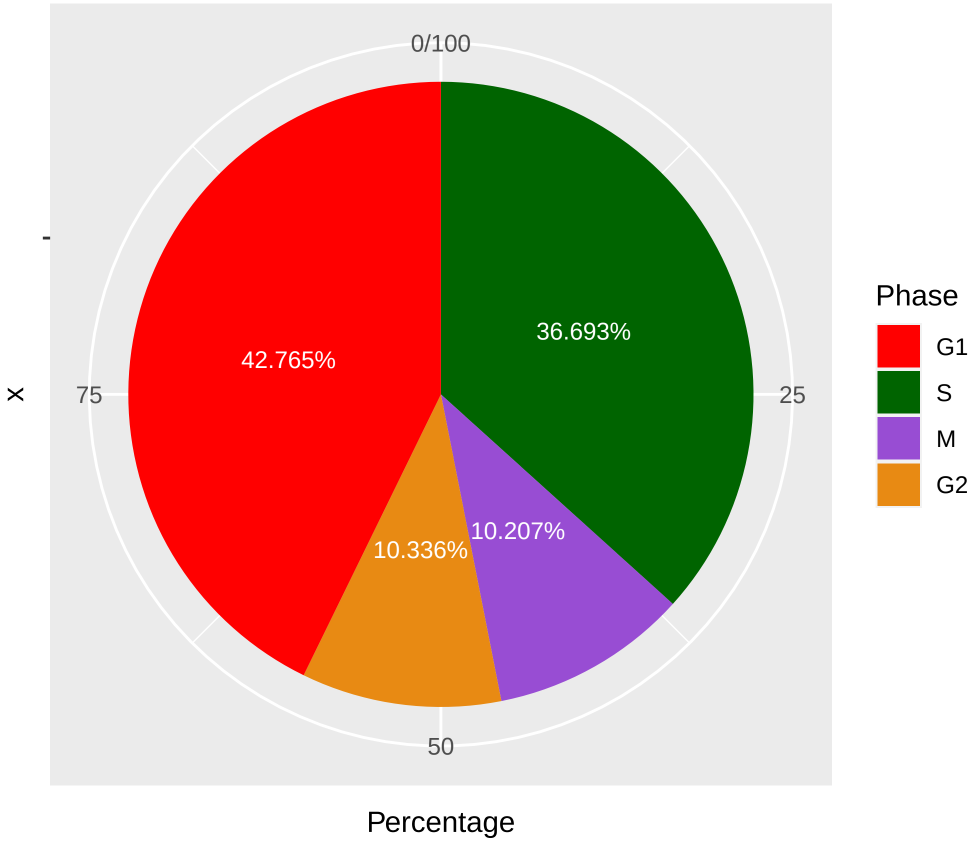
0/100
25
50
75
x
42.765%
36.693%
10.336%
10.207%
Percentage
Phase
G1
S
M
G2
